# Tissue resident CD4+ memory T-cells mark response to immune checkpoint inhibition in high-grade glioma

**DOI:** 10.64898/2026.09.28.754895

**Authors:** Verena Turco, Chin Leng Tan, Dennis Alexander Agardy, Katharina Lindner, Gilbert J. Herrera, Lu Sun, Jakob Rosenbauer, Binghao Zhao, Michael O. Breckwoldt, Lukas Bunse, Robert Prins, Edward W. Green, Michael Platten, Theresa Bunse

## Abstract

**Background:** Immune checkpoint inhibitors (ICI) are efficacious in many solid tumors, but response in glioma is restricted to a small subgroup. The determinants of response and resistance to ICI remain poorly understood.

**Methods:** Here we exploit a syngeneic hypermutated high-grade glioma model with dichotomous response to combined PD-1 and CTLA-4 inhibition to unravel determinants of tumor-infiltrating T-cells driving response. Tumor-infiltrating T-cells from ICI-responsive and non-responsive tumors were analyzed by single-cell RNA and T-cell receptor sequencing and tumor-reactive T-cell receptor clonotypes were functionally validated to characterize their transcriptional phenotypes. We verify our findings in IDH1 wildtype glioblastoma patients treated with neoadjuvant pembrolizumab.

**Results:** ICI response was associated with intratumoral clonal expansion of tumor-reactive cytotoxic T-cells and increased infiltration of CXCR6+ CD4+ tissue resident memory T-cells (Trm). CD4⁺ stem-like memory T-cells in responding tumors demonstrated elevated interferon responses, following trajectories toward clonally expanded Trm, versus trajectories toward exhaustion in non-responsive tumors. In responsive tumors, CD4+ Trm interacted with infiltrating CXCR3+ tumor-reactive and clonally expanded, yet transcriptionally versatile cytotoxic T-cells. Probing the post neoadjuvant ICI high-grade glioma patient tissue dataset, we confirmed increased CXCR6 expression in CD4+ T cells and the association of CD4+ Trm with prolonged overall survival.

**Conclusion:** These findings identify CD4⁺ tissue-resident memory T-cells as determinants of ICI response in IDH1 wildtype high-grade glioma and warrant their further investigation to improve immunotherapy outcomes.

**Key points:**

- Stem-like T cell-derived CD4+ CXCR6+ Trm and tumor-reactive CXCR3+ CD8+ T-cells characterize ICI response in IDH1 wildtype high-grade glioma.
- In IDH1 wildtype glioblastoma patients, CXCR6+ CD4+ Trm associate with prolonged survival after neoadjuvant ICI

**Importance of the study:** We show that CD4+ tissue resident memory cells (Trm) interact with clonal cytotoxic T-cells and are associated with IDH1 wildtype high grade glioma response to ICI and prolonged survival in a syngeneic tumor model and patients treated with neoadjuvant pembrolizumab. Our findings highlight the predictive potential of this specific T cell population, and warrant investigation of CD4+ Trm-targeted combined immunotherapy.

## Introduction

Immune checkpoint inhibition (ICI) targeting immune-inhibitory molecules such as cytotoxic T-lymphocyte-associated antigen-4 (CTLA-4) and programmed cell death protein-1 (PD-1) reinvigorates tumor-reactive T-cells and has demonstrated clinical efficacy in many solid tumors. Yet response remains heterogeneous and is observed only in some patients^1^. Tumor-inherent response determinants are hypothesized to include high tumor mutational burden (TMB), providing neoantigens which are recognized by unleashed neoepitope-specific T-cells^2^ and expression of checkpoint ligands in tumor cells and their microenvironment.

Glioblastomas are highly malignant primary brain tumors with poor prognosis. Several therapeutic approaches including targeted therapy and immunotherapy have not sufficiently improved clinical outcome^3^. Immunologically, glioblastoma is a cold tumor, making immunotherapeutic treatment challenging^4^, although preclinical and early clinical trials suggest that clinical response is possible but restricted to patient subpopulations^5,6^. Resistance to ICI has been attributed to low amounts of tumor-infiltrating T-cells (TILs), an immunosuppressive tumor microenvironment (TME)^7^ and lack of immunogenic neoepitopes ^8^. Nevertheless, anti-PD-1 antibodies induce intratumoral immune responses in preclinical models and in some patients in early clinical trials^6,9,10^, indicating immune recognition. Yet, determinants of ICI response and resistance in isocitratedehaydrogenase (IDH) wt high-grade glioma, especially acquired mechanisms during therapy, remain poorly understood^11^.

To study molecular and cellular response patterns, the immunocompetent syngeneic hypermutated orthotopic IDH1 wt high-grade glioma model GL261 has been comprehensively characterized and is widely used^9,12,13^. We have previously shown reproducibly dichotomous responses to dual ICI of CTLA-4 and PD-1 in this model^9^. Here, response rate was approx. 50% without sex or environmental factors as potential response determinants. This suggests further determinants beyond innate environmental and tumor inherent genetic differences which most likely also apply to patient tumors.

Here, we investigated intratumoral T-cell determinants of response to ICI in the genetically defined GL261 glioma model. Using combined single-cell RNA and T-cell receptor sequencing, we compared tumor-infiltrating tumor-reactive and bystander T-cells from responsive and resistant tumors and validated our findings in clinical data.

## Material and Methods

### Mice

C57Bl/6J mice were used. All animal protocols were performed in compliance to the laboratory animal research guidelines and were approved by the governmental authorities (animal protocols: G-27/17, G-130/23, Regional Administrative Authority Karlsruhe, Germany).

### Tumor cell inoculation and ICI treatment

10^5^ GL261 murine glioblastoma cells were implanted into the right hemisphere. On day 13, allocation to treatment groups was performed based on MRI-based tumor volume. On days 13, 16, and 19, mice were treated with anti-CTLA-4 (9D9) and anti-PD-1 (RMP1-14), or equivalent doses of isotype control antibodies (MCP-11 and 2A3) by intraperitoneal injection.

### Peptide vaccination

C57Bl/6J mice were immunized by subcutaneous injection of 100 µg peptide in Montanide-ISA51 emulsion as described previously^14^. Control mice received emulsion without peptides. Mice were boosted on day 10 and splenocytes were isolated on day 21.

### MR imaging and tumor response criteria

Tumor-bearing animals were imaged using a 9.4 Tesla horizontal bore small animal NMR scanner. Tumor volumes were calculated blinded based on T2 weighted images. Classification of response and non-response was done blinded based on RANO criteria as previously established (*10*).

### Isolation of murine tumor-infiltrating lymphocytes

Mice were cardially perfused and tumor-bearing hemisphere was excised and enzymatically digested with liberase. Myelin was removed by gradient centrifugation.

### Patient Treatment, Tumor Digestion and Isolation of Immune Cells

As described previously^15^, recurrent GBM patients were treated with standard of care therapies, with some receiving off-label, off-trial neoadjuvant pembrolizumab. All patients provided written informed consent. This study was conducted in accordance with the Declaration of Helsinki, and approved by an institutional review board (UCLA Medical Institutional Review Board 2, IRB#10-000655-AM-00059). CD45+ immune cells from a tumor tissue piece were isolated by magnetic bead positive selection.

### TCR cloning

The variable chain of selected murine TCRs was synthesized by Eurofins and cloned into the SMAR-v5 vector using the Golden Gate cloning system and NEB 5alpha competent *E. coli*.

### Generation of TCR RNA

TCR DNA was amplified and a T7 promotor was added. DNA was cleaned and RNA was generated by in vitro transcription using T7 mScript™ Standard mRNA Production System, following the manufacturer’s instructions.

### Isolation of human peripheral blood monocytic cells

Human peripheral blood monocytic cells (PBMCs) were isolated from research-only buffy-coats from healthy donors provided by Blutspendezentrale IKTZ Heidelberg by density gradient centrifugation.

### Rapid expansion of PBMCs

Expansion of PBMC for TCR electroporation was done using irradiated feeder PBMC from three donors and stimulation with human anti-CD3 antibody (OKT-3) and human IL-2, according to the rapid expansion protocol (*97*) (REP) as described previously^16^.

### TCR electroporation

REP-PBMCs were electroporated with TCR-RNA using the 4D Nucleofector (Lonza) according to the manufacturer’s instructions.

### TCR testing

Target tumor cell lines GL261, GL261-OVA I, and BOK were pre-treated with IFN-gamma. TCR-electroporated T (TCR-T) cells and tumor cells were co-incubated at an effector : target ratio of 2:1. Anti-human CD107a antibody was added and incubated for 1 h.

### Flow cytometry

For intracellular cytokine detection, cells were incubated with Brefeldin A or GolgiStop and GolgiPlug. Fixation and permeabilization were performed after blocking and surface staining, followed by intracellular staining. Data were acquired using the Lyric Flow Cytometer (BD Biosciences), or Canto II System (BD Biosciences), and analyzed with FlowJo software (versions 9 or 10). Lymphocytes were subjected to fluorescence-activated cell sorting on FACS Aria II (BD Biosciences; Germany). Alive single CD45high CD3+ cells were sorted from GL261 tumors and used for single cell sequencing.

### Single-cell RNA sequencing (scRNAseq)

#### Murine samples

scRNAseq of sorted CD45+CD3+ T-cells from ICI R and NR murine tumors was performed using 10x Genomics V2 5’ Kit according to manufacturer’s protocol and sequenced on Illumina HiSeq4000. Data was aligned using cellranger (v5.1.2) and analyzed using Seurat (v4.2.0). For trajectory analysis, Monocle was used to calculate the pseudotime and projected onto the UMAP for visualization using Tcf7 cluster as root node.

#### Human samples

scRNAseq was performed according to 10X Genomics 3’ or 5’ gene expression manufacturer’s protocols and sequenced on NovaSeq 6000. Likewise, Data was aligned using cellranger (v7.0.1) and analyzed using Seurat (v4.2.0). For survival analysis, the cohort of neoadjuvant pembrolizumab-treated GBM patients was stratified by their median cell type density value, and survival probabilities were calculated (Kaplan-Meier estimator).

### Whole exome sequencing

Whole exome sequencing (WES) of GL261 tumor tissue from ICI responder and non-responder mice was conducted as described previously^9^.

### Neoepitope MHC-binding prediction

MHC binding prediction of potential GL261-associated neo-epitopes identified by WES was performed using NetMHC 4.0 ^17^ (MHC class I, H2-D^b^ and H2-K^b^), and NetMHCII

2.3^18^ (MHC class II, H2-IA^b^).

### ELISpot (Enzyme-linked-immuno-Spot)

Splenocytes from vaccinated mice were analyzed by IFN-gamma ELISpot as described previously^14^. In some cases, MHC presentation was blocked using blocking antibodies (MHC class I, 28-8-6, and MHC class II, M5/114.15.2). Three technical and at least three biological replicates were used.

## Results

### Activated cytotoxic T-cells of transcriptomic plasticity are enriched in experimental high-grade glioma responding to ICI

Based on heterogenic responses to ICI in glioblastoma patients and in the preclinical syngeneic high-grade glioma model, which can hence not be attributed to inherent genetic differences, we designed this study to evaluate acquired T-cellular response and resistance determinants in the GL261 tumor microenvironment. Although we previously observed differences in glioblastoma-infiltrating myeloid cells between responding (R) and resistant (non-responding, NR) tumors^9^ the ultimate effector cells mediating ICI response are T-cells ^9^. Here, we aimed to identify T-cellular transcriptional programs of response to dual ICI targeting PD-1 and CTLA-4. To this end, we applied dual ICI to GL261-bearing mice as described previously^9^, which showed response rates of approx. 50% of mice defined by preclinical magnetic resonance imaging (MRI)-criteria adopted from clinically established Immunotherapy Response Assessment in Neuro-Oncology (iRANO) criteria^19^ (Fig. 1a-c; Suppl. Fig. 1a-d), and performed combined scRNA- and VDJ-seq on sorted CD45+ CD3+ TILs from R and NR mice to characterize their transcriptomes and TCRs, respectively (Fig. 1d, Suppl. Fig. 1e). Single-cell transcriptomic profiles for a total of 39,178 cells with paired VDJ sequencing were obtained from ICI-treated mice (ICI,R n=5 mice and ICI,NR n=5 mice).

**Figure 1:**
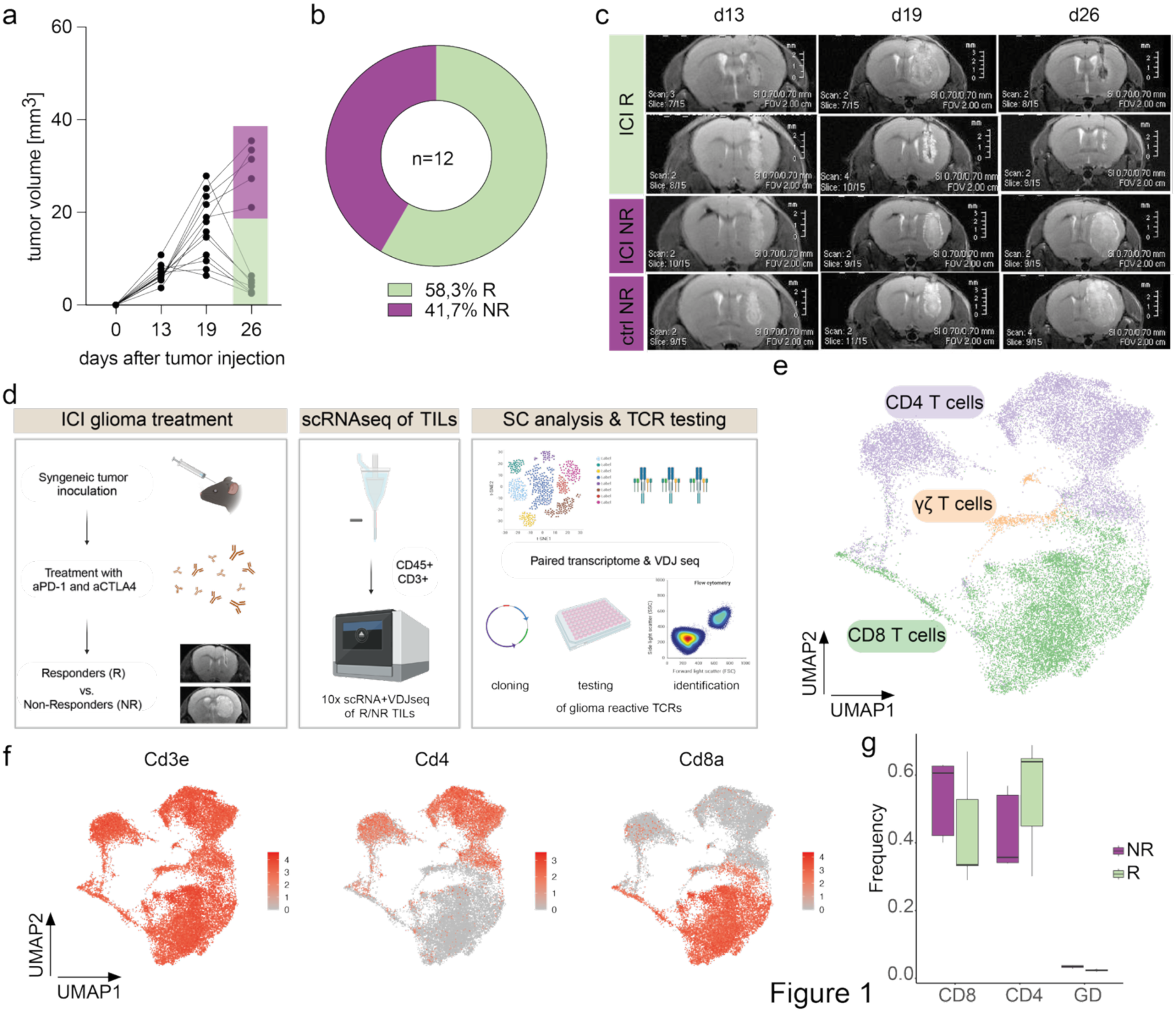
Combined transcriptome and VDJ single cell profiling of GL261 TILs in ICI R and NR mice. **a-d**, C57Bl/6J mice were treated repeatedly with anti-PD-1 and anti-CTLA-4 (ICI) after intracranial GL261 tumor inoculation. Single Cell RNA and VDJ sequencing of tumor-infiltrating T-cells (TILs) of ICI responders (R) and non-responders (NR) was performed on d27. **a**, Quantified tumor growth in ICI treated mice assessed by MRI. Purple, ICI NR; light green, ICI R. **b**, Donut plot of R and NR frequencies based on iRANO criteria. N(ICI R, light green) = 7, n(ICI NR, purple) = 5. **c**, Representative MR images of ICI R (light green) and NR (purple) mice, and vehicle treated control (ctrl, purple). **d**, Workflow for treatment, sample processing, scRNA-seq analysis and TCR testing. **e**, Uniform manifold approximation and projection (UMAP) of CD4+, CD8+, and γδ T-cells within CD45+ CD3+ T-cells obtained from ICI R (n=5) and NR (n=5) mice. n=25387 cells. **f**, Cd3e, Cd4, and Cd8a expression levels across all cells from (e). **g**, Quantified frequencies of CD4+, CD8+, and γδ T-cells within CD45+ CD3+ T-cells from (e). See also Suppl. Fig. 1.

First, we employed unsupervised clustering of CD3+ TILs to segregate CD4+ and CD8+ T-cells for separate comprehensive analysis of both T-cell effector arms (Fig. 1e,f). No significant differences in the ratio of CD4+ and CD8+ T-cell frequencies could be observed between R and NR tumors (Fig. 1g).

To dissect cytotoxic T-cell transcriptomes, we performed unsupervised clustering using louvain algorithm and visualized with uniform manifold approximation and projection (UMAP) of CD3+ CD8+ TILs (n=12,187), identifying 10 distinct transcriptional clusters (Fig. 2a). Differential gene expression (DGE) analysis was performed and used to annotate clusters based on canonical markers^20,21^ (Suppl. Fig. 2a). Phenotypic T-cell states ranged from six active CD8+ TILs (cytotoxic, *Grmc;* activated, *Itgb1;* metabolically active, *Srm;* interferon response, *Isg15;* infiltrating, *Cxcr3;* effector, *Ccl3*) to resident-memory (*Il7r*), terminally exhausted (*Tox*), progenitor-exhausted stem cell-like memory (*Tcf7*) phenotypes, and proliferating CD8+ TILs (*Mki67*). Comparing the CD8+ TIL composition, ICI R tumors were enriched in *Cxcr3*+ CD8+ T-cells (Fig. 2b, Suppl. Fig. 2b,c). These recently infiltrating T-cells have been shown to accumulate in perivascular spaces where they encounter CXCR3 ligands primarily on endothelial cells^22^, suggesting that homing and recruitment of effector TILs to the tumor microenvironment is maintained in R tumors. Moreover, ICI R tumors exhibited a significantly higher frequency of activated CD29+ (*Itgb1*+) CD8+ T-cells compared to NR tumors (Fig. 2b, Suppl. Fig. 2b,c), Human CD29+ CD8+ T-cells have been described to have a high potency of producing IFN-γ and cytotoxic molecules thus enhanced killing capacity^23^. We previously showed that CD29 is upregulated upon antigen-specific activation and required for effector functions, while engagement with its ligand osteopontin enhances expression of the exhaustion marker Tox^24^. The frequency of terminally exhausted *Tcf7-Tox+* CD8+ T-cells, however, was not significantly increased in R compared to NR TILs (Fig. 2b, Suppl. Fig. 2b,c), suggesting ICI-dependent control of the terminal exhaustion susceptibility of cytotoxic CD29+ T-cells. Accordingly, we did not observe an increase in global CD8+ T-cell exhaustion (Suppl. Fig. 2d), although this is regularly observed following local antigen recognition and clonal T-cell expansion. Similarly, we did not detect an increase in global cytotoxicity in CD8+ T-cells from ICI R tumors Suppl. Fig. 2e).

**Figure 2:**
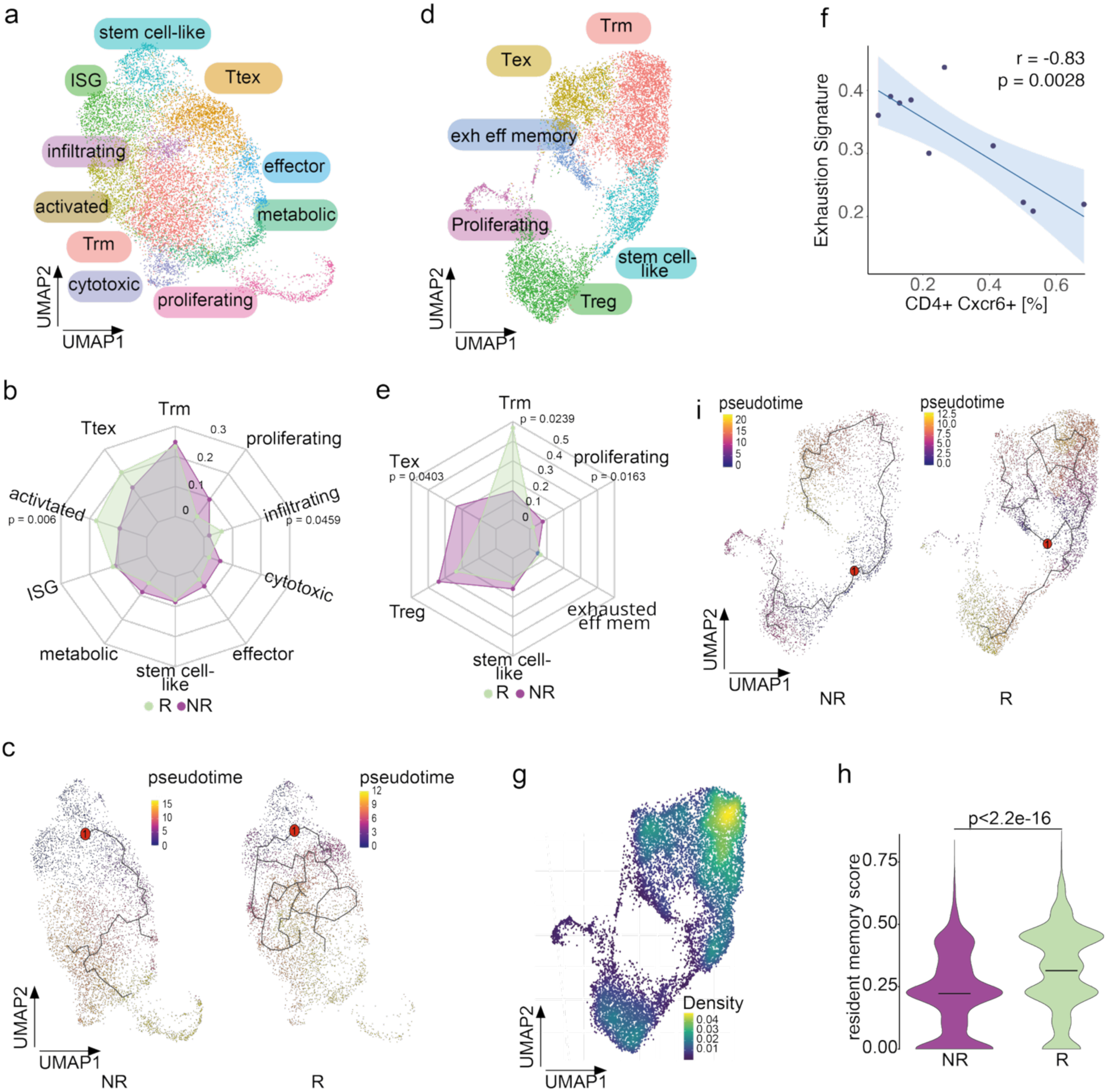
Transcriptomic plasticity of cytotoxic T-cells and CD4+ tissue resident memory T-cells are associated with preclinical ICI response. **a,** UMAP of CD45+ CD3+ CD8+ T-cell clusters obtained from ICI R (n=5) and NR (n=5) mice. Phenotypic clusters are represented in distinct colors. n = 12,187 cells. **b,** Radar plot depicting CD8+ T-cell cluster distribution in ICI R (light green) and NR (purple). Significant p values are depicted. T-test with Holm correction. **c,** CD8+ T-cell pseudotime analysis using Monocle in ICI R and NR. *Tcf7*+ stem cell-like cluster was chosen as the root node (red dot). **d,** UMAP of CD45+ CD3+ CD4+ T-cell clusters obtained from ICI R (n=5) and NR (n=5) mice. Phenotypic clusters are represented in distinct colors. n = 11,795 cells. **e,** Radar plot depicting CD4+ T-cell cluster distribution in ICI R (light green) and NR (purple). Significant p values are depicted. T-test with Holm correction. **f,** Correlation of exhaustion signature (*Havcr, Lag3, Ctla4, Pdcd1, Tigit, Tox*) expression and *Cxcr6* expression levels in CD4+ T-cells from ICI R and NR (n = 10) mice. Pearson correlation. **g,** Feature UMAP depicting Trm score (*Cxcr6*, *Il7r*, *Itga1*, *Dusp6*) density across CD4+ T-cell clusters from d. **h,** Resident memory score expression levels in ICI NR and R CD4+ T-cells. Score as in (g). n = 7143 (R) and 4652 (NR) cells. Welch t-test. **i,** CD4+ T-cell pseudotime analysis using Monocle in ICI R and NR mice. *Tcf7*+ stem cell-like cluster was chosen as the root node (red dot). See also Suppl. Fig. 2,3.

In summary, responding tumors contained an increased proportion of *Cxcr3+* recently infiltrating and *Itgb1+* highly activated cytotoxic T-cells without a secondary increase in terminal exhaustion.

Response to ICI has been associated with the reprogramming of progenitor exhausted CD8+ *Tcf7*+ T-cells^25^. Therefore, we performed pseudotime analysis with the progenitor exhausted CD8+ *Tcf7*+ cluster as root node. *Tcf7* encodes the transcription factor T-cell factor 1 (TCF1), which characterizes a unique stem cell-like differentiation state that gives rise to central memory cells^26^. In ICI R, we found multiple trajectories toward and around activated recently infiltrating and resident memory CD8+ T-cell clusters, while their ICI NR counterparts followed a clearer trajectory along the terminally exhausted cluster (Figure 2c). These results indicate a transcriptomic plasticity in cytotoxic T-cells exerting response to ICI. To confirm that this plasticity is a consequence of defined cellular trajectories but not transcriptional alterations in the root node, we performed DGE analysis of the progenitor exhausted *Tcf7*+ CD8+ T-cell clusters. Supporting our hypothesis, this revealed only 3 DEG between *Tcf7*+ CD8+ T-cells from R and NR tumors (Suppl. Fig. 2f). Taken together, activated and recently infiltrating cytotoxic T-cells are enriched in ICI responding tumors because of transcriptomic plasticity originating from progenitor exhausted T-cells, while global cytotoxic T-cell exhaustion remains unaffected.

### Tissue resident memory CD4+ T-cells are associated with ICI response of experimental glioblastoma

In previous work in human and experimental high-grade glioma, we found a unique CD4+ T-cell-MHC class II-dependent exhaustion susceptibility of CD29+ tumor-reactive T-cells^24^ and T helper cells to be equally important for response to ICI^9^. Hence, we hypothesized that control of global exhaustion despite a higher abundance of CD29+ cytotoxic T-cells is controlled by CD4+ T-cells. Thus, we analyzed their transcriptomic landscape in R and NR tumors. Unsupervised clustering of 11,795 CD3+ CD4+ TILs identified six distinct transcriptional clusters, namely a *Tcf7+* stem cell-like memory cluster, similar to the respective cytotoxic T-cell cluster, exhausted (*Havcr*), exhausted effector memory (*Tox2*), tissue-resident memory (*Cxcr6*), proliferating (*Mki67*), and regulatory T-cell (T_reg_, *Foxp3*) clusters (Fig. 2d, Suppl Fig. 3a).

To define differences in CD4+ T-cell transcriptomes between ICI responding and resistant tumors, we first compared cluster frequencies. In line with our studies, ICI resistant tumors harbored an enhanced frequency of exhausted *Havcr* expressing CD4+ T-cells (Fig. 2e; Suppl. Fig. 3b-d), indicating failed reinvigoration under PD-1 and CTLA4 blockade. *Havcr* encodes TIM3, which marks the most dysfunctional subgroup among tumor-infiltrating PD1+ cytotoxic and CD4+ T-cells^24^ and can be upregulated upon PD-1 blockade in a compensatory resistance mechanism^27^. Accordingly, in preclinical and patient studies, dual checkpoint inhibition using synergistic TIM3 and PD-1 blockade improved response^28^ (NCT02817633), underlying the relevance of *Tim3*-expressing T-cells for resistance to PD-1 ICI.

Interestingly, *Cxcr6*-expressing tissue-resident memory (Trm) CD4+ T-cells were significantly enriched in ICI R tumors (Fig. 2e; Suppl. Fig. 3b-d). The chemokine receptor CXCR6 supports Trm functionality and tissue retainment via binding to its ligand CXCL16 on myeloid cells^29^. Hence, in contrast to recently infiltrating *Cxcr3*+ cytotoxic T-cells, ICI R tumors contained more tissue-retained parenchymal CD4+ T-cells. Transcriptionally, *Cxcr6* levels correlated negatively with an exhaustion signature in CD4+ T-cells (Figure 2f), supporting their association with a memory phenotype. Further, we calculated a tissue resident memory score, confirming its highest expression in the Trm cluster (Fig. 2g) and elevated expression in CD4+ T-cells from responding tumors (Fig. 2h).

To probe the hypothesis that enriched activated parenchymal Trm CD4+ T-cells in ICI R tumors arise from trajectories of stem-like *Tcf7*+ CD4+ T-cells, we again performed pseudotime analysis with the stem cell-like cluster as root node. Indeed, progenitor exhausted *Tcf7+* cells showed trajectories towards and within the Trm cluster and towards the Treg cluster in responding tumors, while those from NR tumors showed clear trajectories towards exhausted and Treg clusters (Fig. 2i). Since CD4+ T-cells strongly interact with the myeloid-rich glioma microenvironment, we hypothesized that the CD4+ stem cell-like memory T-cell cluster in ICI R tumors specifically harbors transcriptional features determining cellular trajectories. Remarkably, DGE analysis revealed that these T-cells from NR tumors expressed elevated levels of activation markers such as *Cd74, Gzma* and *Grmb*, and Treg-specific genes *Foxp3* and *Lrcc32*^30^ (Suppl. Fig. 3e), likely making them prone to Treg differentiation and exhaustion. Conversely, elevated expression of only two genes could be detected in CD4+ stem cell-like memory T-cells from R tumors, incl. the interferon response gene *Ifitm10*, which has been implicated in T-cell antitumor immune function^31^. Accordingly, gene ontology analysis revealed enrichment of interferon-response pathways in these cells. (Suppl. Fig.3f).

Collectively, these data suggest that ICI response of experimental high-grade glioma is associated with an enrichment of intratumoral CD4+ resident memory T-cells directly or indirectly recruiting *Cxcr3*+ recently infiltrated cytotoxic T-cells, potentially controlling their function.

### Tissue-resident memory CD4+ T-cells are clonally expanded, and CD4 T-cells recognize a GL261 neoantigen in ICI responsive tumors

Next, we asked if the higher frequency of CD4+ Trm results from increased infiltration or local antigen-specific expansion. Analysis of αβ-paired VDJ sequences of CD45+ CD3+ TILs revealed lower VDJ diversity in 4/5 (80%) ICI R animals compared to ICI NR mice (Fig. 3a,b), indicating greater global clonality and contraction of the TCR repertoire under successful ICI therapy. This was further emphasized by the scarcity of rare clones in ICI R mice (Fig. 3b). Yet we observed a substantial interindividual heterogeneity of the VDJ repertoire irrespective of response to ICI. Despite the use of a genetically defined tumor model with injection of a cell line into inbred mice, VDJ sequence similarity as estimated by the Morisita-Horn-index indicated no relevant overlap between individual animals (Fig. 3c). Nevertheless, increased CDR3 sequence similarity across all TILs from ICI R tumors (Fig. 3d) suggests recognition of multiple tumor antigens. To link transcriptomic profiles to VDJ clonotypes, we integrated VDJ clonotypes into transcriptomic clusters and superimposed the top 5 clonotypes of all R and NR mice on all respective T-cell clusters (Fig. 3e). Strikingly, 3/5 R tumors displayed a CD4+ T-cell-dominated top VDJ repertoire localized mainly in Trm cluster, whereas in NR mice, high clonality was found in exhausted CD4+ T-cells. In CD8+ T-cells, top clonality was more heterogenous and observed mainly in resident memory, terminally exhausted, and activated clusters, again indicating a transcriptomic plasticity of clonally expanded CD8+ T-cells, while ICI R is characterized by clonal expansion of CD4+ Trm.

**Figure 3:**
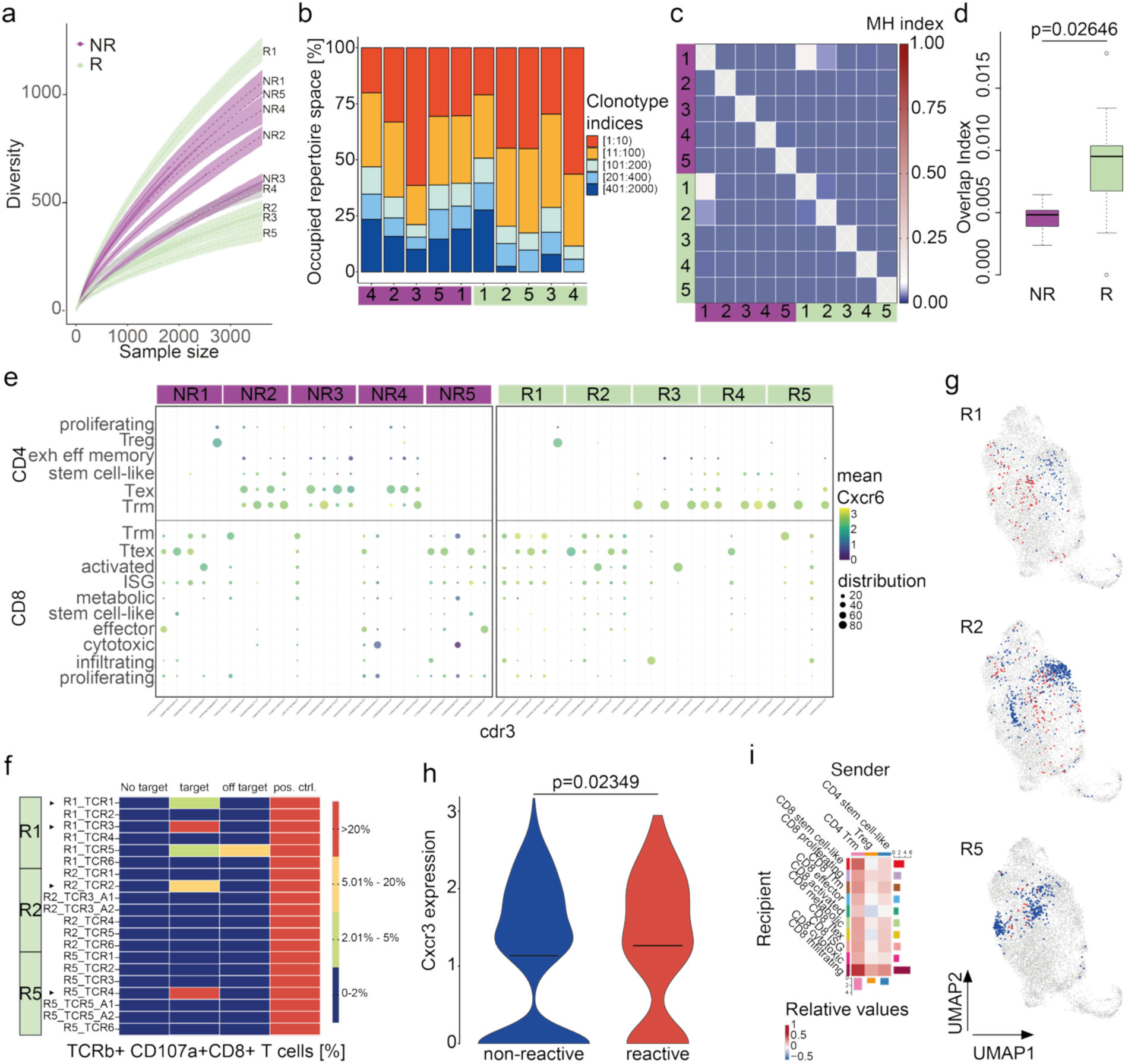
CD4+ Trm cells are clonally expanded and tumor-reactive transcriptomically plastic cytotoxic T-cells express high levels of *Cxcr3* in preclinical high-grade glioma. **a,** TCR diversity in individual ICI R (light green, n = 5) and NR (purple, n = 5) mice. Numbers depict individual mouse identities. **b**, Clonotype distribution of TCRs in individual ICI R and NR mice as in a. purple, NR mice; light green, R mice. Numbers depict individual mouse numbers as in (a). **c**, Morisita-Horn-index comparing individual TCR sequences from ICI R (light green) and NR (purple) mice as in (a). **d**, CDR3 sequence similarity by overlap index in T-cells from ICI R ( n = 5) and NR (n = 5) mice. **e**, Cluster distribution and mean *Cxcr6* expression levels of top five CD4+ (top) and CD8+ (bottom) TCR clones per individual mouse in ICI R (n = 5, light green) and NR (n = 5, purple) mice. Numbers depict individual mouse numbers as in (a). **f**, Heatmap depicting reactivity of tested TCRs (n = 20) from three ICI R (light green) mice. Reactivity was determined by co-culture of TCR-transfected T-cells with GL261 glioma cells (targeT-cell line). Positive flow cytometry staining of TNF-α and/or CD107a on successfully transfected T-cells at a frequency above those co-cultured with an off-targeT-cell line was defined as reactivity. TCR defined as reactive are marked with an arrow. pos. ctrl., positive control, stimulation with CD3/CD28 beads. Numbers depict individual mouse numbers as in (a). **g**, Tested TCRs from (f) and their reactivities superimposed on individual CD8+ T-cell UMAPs from R mice. red, reactive clones; blue, non-reactive clones; grey, non-tested clones. Numbers depict individual mouse numbers as in (a). **h**, *Cxcr3* expression levels of CD8+ T-cells carrying reactive or non-reactive TCRs defined in (f) and as in (g). N(reactive) = 279 cells, n(non-reactive) = 1,579 cells. T-test. **i**, Heatmap depicting relative interaction strength between CD4+ and CD8+ T-cell clusters from ICI R compared to NR. See also Suppl. Fig. 4,5.

GL261 represents a hypermutated high-grade glioma model and neoepitopes are largely recognized by CD4+ T-cells, thus we hypothesized that clonal expansion of CD4+ Trm reflects endogenous neoepitope recognition. With the attempt to identify such neoepitopic antigens, we retrieved point mutations from GL261 tumors ex vivo after ICI by whole exome sequencing (WES) and selected those enriched in resistant tumors but absent in responsive tumors, hypothesizing that tumor cells expressing relevant antigens have been successfully eradicated in R tumors (Suppl. Fig. 4a,b). In silico MHC binding prediction via NetMHC revealed five potential class II-restricted neoepitopes, two of which were predicted mutation-specific (Suppl. Fig. 4c,d). These five potential neoepitopes were tested for immunogenicity by peptide vaccination of C57BL/6 mice, and one proved immunogenic, eliciting mutation-specific IFN-γ T-cell responses after vaccination. (Suppl. Fig. 4e,f). Blocking antigen presentation on MHC class I and II molecules during ex vivo recall, we validated MHC class II restriction, confirming CD4+ T-cell recognition and supporting our hypothesis (Suppl. Fig. 4g,h).

### GL261-reactive cytotoxic T-cells express high levels of *Cxcr3* and are transcriptionally plastic

We found that ICI response in experimental glioblastoma is associated with an enrichment of intratumoral clonally expanded CD4+ Trm which may recruit cytotoxic T-cells. Since the latter can be exploited for TCR-engineered T-cell therapy, we selected putative tumor-reactive TCRs based on frequency and tested the top 6-7 clonally expanded CD8+ TCRs of three individual ICI-responsive mice by overexpressing them in rapidly expanded PBMCs by RNA-based electroporation and analyzing reactivity against the target cell line GL261in a short-term *in vitro* co-culture by flow cytometry (Suppl. Fig. 5a-c). Among 20 TCRs tested, we extracted four reactive TCRs eliciting variable anti-tumor responsiveness despite comparable TCR expression levels after electroporation, with few exceptions (Fig. 3f, Suppl. Fig. 5d,e), implying that ICI response-associated CD8+ TCRs have varied affinity for GL261 tumor epitopes.

To define transcriptomic signatures of glioma-reactive T-cells which may determine ICI response, we superimposed VDJ sequences of tumor-reactive T-cells on transcriptomic clusters of individual mice. Reactive clonotypes were distributed across multiple T-cell clusters, both interindividually and individually (Fig. 3g), demonstrating that T-cell clones co-exist in multiple transcriptional states and confirming their transcriptomic plasticity. Nevertheless, reactive CD8+ T-cells expressed increased *Cxcr3* levels, indicating their recent infiltration (Fig. 3h). Finally, to understand how clonal CD4+ Trm affect CD8+ T-cell antitumor reactivity, we hypothesized that parenchymal CD4+ Trm recruit recently infiltrated CD8+ T-cells in R tumors. Therefore, we performed receptor-ligand analyses on recombined CD4+ and CD8+ T-cell transcriptomes. As CD4+ T-cells trajected mainly from a stem cell-like phenotype to Treg or Trm, we analyzed interactions of these transcriptomic clusters with CD8+ T-cells. Supporting the importance of CD4+ Trm recruiting active cytotoxic T cells for ICI response, we found the strongest interaction between CD4+ Trm senders and recently infiltrated CD8+ T-cells among all CD8+ recipient clusters in R tumors (Fig. 3i).

In line with our trajectory analyses, this highlights both antigen- and ICI-orchestrated trajectory programs in ICI responsive tumors. In sum we find that infiltrating CD8+ T-cells express tumor-reactive TCRs which co-evolve with clonally expanded, in part neoantigen-reactive CD4+ Trm in ICI-responsive tumors.

### CXCR6 expression in tumor-infiltrating CD4+ Trm cells is associated with clinical response to neoadjuvant checkpoint blockade in high-grade glioma patients

In patients, CXCR6 is associated with prolonged survival in many solid cancers, including melanoma, head and neck, breast, colorectal and ovarian cancer^29,32,33^ Its relevance for ICI response and clinical outcome has recently been suggested in glioblastoma^34^. However, CXCR6 expression has neither been systematically or specifically analyzed post PD-1 blockade in high-grade glioma patients nor contextualized with a specific co-evolution of CD4+ and CD8+ T-cells. Similarly, CD8+ but not CD4+ Trm have been shown to be associated with prolonged survival in cancer patients^35^. Hence, we analyzed *CXCR6* expression in human glioblastoma-infiltrating T-cells using our publicly available scRNA-seq data from patients treated with neoadjuvant pembrolizumab (anti-PD-1), which has shown survival benefit in selected patients with recurrent glioblastoma^6^. Interestingly, *CXCR6* expression was increased in TILs following anti-PD-1 therapy^15^. Re-analysis of single-cell data confirmed *CXCR6* upregulation in pan T-cells after PD-1 blockade (Fig 4a, Suppl. Fig. 6a). We specified these findings in an independent controlled scRNA-Seq dataset of TILs from recurrent isocitrate dehydrogenase wildtype (IDHwt) glioblastoma patients treated with neoadjuvant pembrolizumab (n=21) or untreated (n=21), in which CD4+ T-cell *CXCR6* expression was elevated after PD-1 blockade (Fig. 4b-c, Suppl. Fig. 6b). Of note, we confirmed the presence of myeloid cell clusters expressing the CXCR6 ligand *CXCL16* (Suppl. Fig. 6c-d), which may be responsible for Tcm recruitment into the TME. Interestingly, among four CD4+ T-cell clusters, defined as Treg, central memory T-cells (Tcm), Trm, and stress response T-cells (Tstr), Trm and Treg expressed the highest CXCR6 levels (Fig. 4d). Excluding Tregs in a survival analysis, we found that a high density of *CXCR6*-expressing Trm correlated significantly with prolonged overall survival (OS) among Pembrolizumab-treated patients compared to a low density of these cells (Fig. 4e). In contrast, the association of a recently-infiltrating CXCR3+ CD8+ T-cell cluster density with prolonged OS was insignificant in this patient group (Suppl. Fig. 6f,g), suggesting that other cytotoxic such as progenitor exhausted T-cells are more relevant for response to PD-1 blockade in these patients^25^.

**Figure 4:**
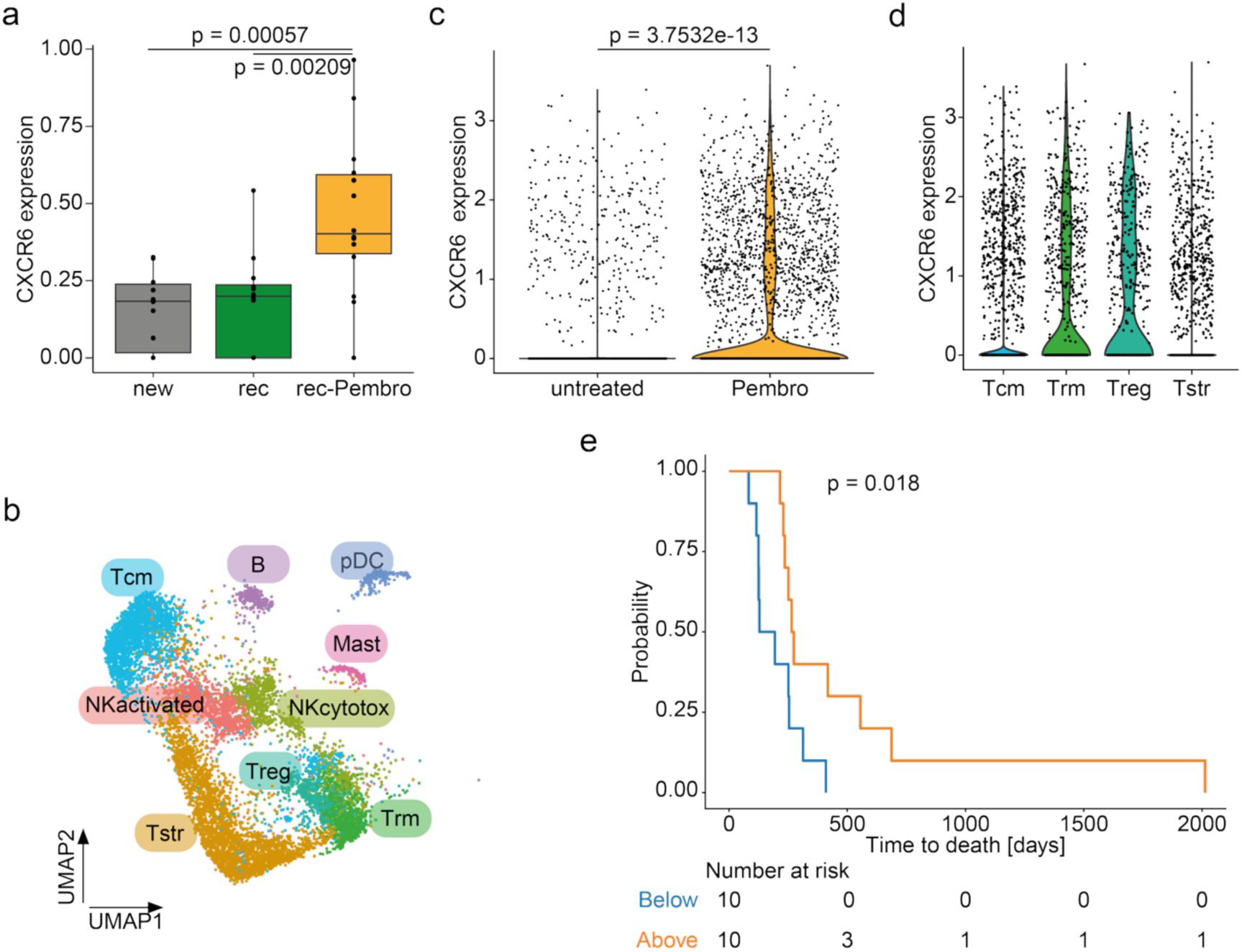
CD4+ *CXCR6*+ Trm cells are associated with response to pembrolizumab in high-grade glioma patients. **a**, *CXCR6* expression levels in bulk T-cells from newly diagnosed (new), recurrent (rec), and pembrolizumab-treated recurrent (rec-Pembro) glioblastoma samples^15^. n(new) = 9, n(rec) = 9, n(rec-Pembro)=13. T-test with Holm correction. **b**, UMAP of *CD8A*- and *CD8B*-lymphoid cells from untreated (n = 15) or neoadjuvant pembrolizumab-treated (Pembro, n = 20) recurrent glioblastoma samples, indicating CD4+ T-cell and other clusters. Phenotypic clusters are represented in distinct colors. N = 7,509 cells. **c**, *CXCR6* expression levels in CD4+ T-cells from untreated (n = 15) or neoadjuvant pembrolizumab-treated (Pembro, n = 20) recurrent glioblastoma samples. n(untreated) = 2,294 cells, n(Pembro) = 5,215 cells. Wilcoxon rank sum test. **d**, *CXCR6* expression levels in CD4+ T-cell clusters. n(Tcm) = 2,459 cells, n(Trm) = 1,012 cells, n(Treg) = 540 cells, n(Tstr) = 3,185 cells. Tcm, central memory T-cells, *CCR7*+; Trm, tissue resident memory T-cells, *CXCR6*+ *FOXP3*-; Treg, regulatory T-cells, *FOXP3*+; Tstr, T-cells with stress response, HSPA1A+. **e**, Kaplan-Meier plot of overall survival probability of glioblastoma patients treated with neoadjuvant pembrolizumab as in b according to Trm density among CD4+ T-cells. Orange, above median Trm density; blue, below median Trm density. N(above) = 10, n(below) = 10. Mantel-Cox log-rank test. See also Suppl. Fig. 6.

Collectively, these data verify the relevance of CD4+ CXCR6+ Trm for the response to ICI in high-grade glioma patients.

Taken together, our findings define the immunological T-cell landscape in response to dual ICI therapy, bridging TCR clonality, phenotypic diversity, and treatment effects. Our data demonstrate the substantial interindividual heterogeneity of intratumoral pan and tumor-specific T-cell clones in an experimental syngeneic inbred mouse glioblastoma model. Importantly, we establish CD4+ Trm as a shared determinant of ICI response in both preclinical and clinical high-grade glioma, which might serve as an early predictive marker for ICI in patients.

## Discussion

Neo-adjuvant ICI has been investigated in high-grade glioma in clinical trials, where it improved survival in a subset of patients^6^. In this study, we exploited dichotomous response to ICI in a syngeneic, hypermutated glioma model to investigate acquired transcriptomic and clonal heterogeneity of tumor-infiltrating T-cell responses at single-cell resolution. In ICI-responding tumors, we found an increased abundance of effector CD8+ T-cells expressing high levels of CD29, encoded by *Itgb1*, which is upregulated after activation and important for effector functions, as well as those that express the chemokine receptor Cxcr3, mediating migration (Fig. 2a-b). CD29+ effector CD8+ T-cells are known for their high cytotoxic capacity and have been associated with favorable prognosis in melanoma^23^. We have previously shown that CD29 is upregulated in cytotoxic T-cells upon antigen encounter and indispensable for their effector function^24^. However, CD29 can have a dual role by engaging in terminal CD8+ T-cell exhaustion in MHC class II-deficient microenvironments due to the lack of CD4+ T-cell activation which leads to expression of the terminal exhaustion marker Tox^24^ ^36^. Importantly, in our dataset, arguing against a stronger terminal exhaustion in NR tumors and for reinvigoration by a third immune checkpoint blocker^9^. The CXCR3– ligand axis plays a critical role for T-cell trafficking and function^37^ ^38^. CXCR3 is typically highly expressed on effector CD8+ T-cells and promotes T-cell migration from the periphery to the tumor microenvironment^39^ or into the brain, e.g. during cerebral infection^40^. CXCR3 facilitates interaction with antigen-presenting cells^41^, thus potentially contributing to anti-tumor response in ICI R mice. Consistent with this, CXCR3-dependent T-cell tumor accumulation is required for successful anti-PD-1 therapy in experimental melanoma^42^ and has been linked to ICI efficacy in metastatic urothelial cancer, suggesting a broader role for CXCR3-mediated homing.

The most prominent changes upon response to ICI were observed within CD4+ T-cells, reflecting the importance of these for ICI response in this model^9^, and supporting the notion that CD4+ T-cell phenotypes determine whether a cytotoxic T-cell response, which may have been initiated also in ICI resistant tumors, becomes effective. ICI-responsive tumors were characterized by a reduced frequency of *Tim-3*-expressing exhausted CD4+ T-cells, indicating insufficient reinvigoration in ICI-resistant tumors (Fig. 2d,e) and underlying the essential role of TIM3 for ICI resistance. It marks the most dysfunctional subgroup among tumor-infiltrating PD1+ cytotoxic and CD4+ T-cells^43^ and can be upregulated upon PD-1 blockade^27^, reflecting acquired resistance.

Remarkably, ICI-responsive tumors harbored a highly increased frequency of CD4+ Trm (Fig. 2e,g,h), which was due to differentiation from stem cell-like memory T-cells, while these trajected toward exhausted or regulatory phenotypes in resistant tumors (Fig. 2i), likely determined by expression of immunosuppressive and Treg-associated genes in stem-like T-cells in NR tumors (Suppl. Fig. 3e).

The CD4+ Trm cluster was characterized by several tissue residency markers, including *Cxcr6* and *Il7r*, whose combined expression was also upregulated in ICI R TILs (Fig. 2g,h). CXCR6 is crucial for functionality, survival and tissue retainment^29,44^. Accordingly, CXCR6-expressing T-cells have been associated with effective cytotoxic immune responses in human cancer such as non-small cell lung cancer^45^. In preclinical models, *Cxcr6* expression was increased in infiltrating CD8+ T-cells after anti-PD-1 treatment, and its deficiency abrogated cytotoxic T-cell anti-tumor functionality^32^, while its overexpression by transferred CAR T-cells enhanced their intratumoral accumulation and survival of mice^46^. However, its role in CD4+ T-cells has not been described previously. IL7R is essential for IL-7–mediated T-cell survival, homeostasis, and memory formation and is expressed in most mature CD4+ T-cell subsets except Tregs^47^. In CD4+T-cells, high *Il7r* expression is associated with an antigen-specific central memory phenotype and favorable persistence, proliferation and antitumor capacity in the context of immunotherapy and adoptive T-cell therapy^48,49^.

CD8+ Trm cells have been shown to play an important role for enhanced survival in cancer; however, the underlying mechanisms remain unclear^35^. Tumor-resident memory T-cell subsets have been demonstrated to regulate responses to anti-PD-1 and anti-CTLA-4 cancer immunotherapies in melanoma and lung cancer^50^. Similarly, in multiple sclerosis (MS) patients, CD8+ Trm cells have been found in white matter lesions^51^, whereas the only evidence for a role of CD4+ Trm cells in CNS immune responses comes from progressive MS patients and the MS model experimental autoimmune encephalomyelitis (EAE), where CD4+ proinflammatory Trm seem to significantly contribute to pathogenesis^52^. Considering these studies, we provide first evidence that CD4+ Trm cells contribute to immunotherapy responses against primary brain tumors.

In responding tumors, Tcf7+ stem–like CD4+ T-cells showed only marginal upregulation of few genes, mainly related to IFN responses. In contrast, NR Tcf7+ stem cell-like CD4+ T-cells displayed enhanced expression of the Treg gene *Lrrc32, which* encodes the surface protein Glycoprotein A Repetitions Predominant (GARP). GARP tethers transforming growth factor β (TGF-β) to the cell surface, thereby rendering it latent^30^. TGF-β is activated when it is released from this complex^53^. In turn, TGF-β is required for the development of CD8+ Trm. While Treg can provide TGF-β in this process^54^, it remains speculative if an enhanced *Lrrc32* expression and hence surface-tethered TGF-β by Treg – or by stem cell-like CD4+ T-cells on their trajectory towards Treg cells – may inhibit resident memory T-cell formation. If so, increased GARP levels in Tcf7+ stem cell-like NR T-cells may have a dual role in determining their trajectory by (i) skewing these cells towards a Treg phenotype, and (ii) preventing TGF-β-mediated development of Trm.

Given the significantly higher abundance of *Cxcr6+* CD4+ Trm in responding experimental glioma, we verified CXCR6 prognostic relevance in high-grade glioma patients treated with neoadjuvant pembrolizumab. CXCR6 is part of two pan-cancer gene signatures predictive of anti-PD-1 response^55,56^. Similarly, we found higher *CXCR6* expression in pre-treatment tumor-infiltrating T-cells from pembrolizumab-responsive high-grade glioma patients. A high proportion of *CXCR6*-expressing T helper cells was associated with prolonged overall survival in these patients. Interestingly, we found high CXCR6 expression in Treg. Similarly, in a melanoma model, CXCR6 could be detected in infiltrating Treg; however, Treg contribution to CXCR6 expression in the entire TME was marginal^29^. This argues for a comprehensive analysis when using CXCR6 as a predictive marker in high-grade glioma.

Biologically, the CXCR6–CXCL16 axis supports T-cell effector function via NF-κB– dependent cytokine production, and its relevance is validated by the robust expression of CXCL16 in myeloid cells, specifically in conventional DC, within our glioblastoma dataset (Suppl. Fig. 6d).

Our analysis of scVDJ repertoires revealed decreased TCR diversity and increased clonality in ICI responding tumors, consistent with antigen-driven clonal expansion. Despite enhanced CDR3 sequence similarity in responding tumors (Fig. 3d)^57,58^, the post-ICI intratumoral TCR repertoire showed remarkable interindividual heterogeneity, in both responding and resistant tumors, although in a syngeneic, genetically defined inbred mouse strain. This suggests stochastic TCR clonal expansion upon ICI, potentially modulated by antigen availability, tumor microenvironmental cues, and immunoediting during ICI treatment^59^. In fact, the theoretical maximum TCR diversity in the mouse is as large as 10^15^, while the spleen of an immunologically naïve mouse harbors only 2×10^6^ clones^60^. Understanding determinants of TCR heterogeneity in standardized preclinical models can provide valuable insights in immunological mechanisms underlying ICI response.

Integration of transcriptomic and VDJ data (Fig. 3e) revealed that in R tumors, top CD4+ TCR clones were Trm, while in NR tumors, top, putatively antigen-specific, CD4+ clones were exhausted. This suggests that antigen-specific Trm determine ICI response. Within the CD8+ T-cell compartment, however, we identified substantial heterogeneity in the transcriptomic state of top TCR clonotypes. In an antigen-agnostic approach, we found that functionally validated tumor-reactive, clonally expanded TCRs were distributed across several CD8⁺ T-cell states. This suggests that ICI response is orchestrated by coordinated yet heterogeneous cytotoxic T-cell programs, and that clonal expansion may be a surrogate for antitumor-specificity, but does not predict functional antitumor activity. Nevertheless, the increased *Cxcr3* expression by reactive CD8+ T-cells (Fig. 3h) supports the hypothesis that antigen-specific CD4+ Trm recruit antigen-specific cytotoxic T-cells in an anti-tumor immunological triad that involves CXCL16+ myeloid cells.

In conclusion, we identify CD4+ Trm and their interaction with CXCR3+ cytotoxic T-cells as key determinants of ICI response, potentially serving as a predictive biomarker in future clinical trials. Strategies to enhance response to ICI in combinatorial therapeutic settings might involve *CXCR3* overexpression in T-cell products as well as approaches enabling targeted reinvigoration of CD4+ Trm cells.

## Supporting information

Supplementary data incl methods with method tables and supplementary figures

## Required statements

### Ethics

All animal protocols were performed in compliance to the laboratory animal research guidelines and were approved by the governmental authorities (animal protocols: G-27/17, G-130/23, Regional Administrative Authority Karlsruhe, Germany).

### Authorship

V.T. designed and performed in vivo and in vitro experiments and analyzed data. C.L.T. designed experiments and analyzed murine and human data. K.L. and D.A.A. performed in vitro experiments. G.J.H., L.S., J.R., and B.Z. analyzed human data. M.O.B. analyzed and interpreted MRI measurements. L.B. and E.W.G. interpreted data. R.P. provided and interpreted data. M.P. and T.B. conceptualized the study and interpreted data. V.T., C.L.T. and T.B. wrote the paper with input from all co-authors.

### Data Availability

The data generated in this study will be available within the article and its supplementary files. RNA sequencing data will be made available upon publication. All other raw data will be made available upon reasonable request to the corresponding author.

## Acknowledgements

We are grateful for the patients and their relatives who participated in the clinical trial. We acknowledge the support from the Core Facility for Flow Cytometry (Dr. Steffen Schmidt) and the Single Cell Open Lab (Dr. Jan-Philipp Mallm) at the German Cancer Center. We acknowledge the data storage service SDS@hd supported by the Ministry of Science, Research, and the Arts Baden-Württemberg (MWK) and the German Research Foundation (DFG) through grant INST 35/1314-1 FUGGand INST 35/1503-1 FUGG. We thank Anna von Landenberg, Kristine Jähne, Gordon Haltenhof and Manuel Fischer for excellent technical support. V.T. received fellowships from the German Academic Scholarship Foundation (SDV) and the Mildred-Scheel doctoral program of the German Cancer Aid. D.A.A. was funded by Deutsche Forschungsgemeinschaft (German Research Foundation, DFG)–project number 259332240, RTG 2099 Hallmarks of Skin Cancer, and by DFG–project ID 404521405, SFB 1389–UNITE Glioblastoma, WP B01. J.R. was funded by DFG–project ID 404521405, SFB 1389–UNITE Glioblastoma, WP B01. K.L. was funded by the Helmholtz International Graduate School for Cancer Research. M.O.B. was supported by the Emmy Noether program of the German Research Foundation (DFG, BR 6153/1-1), the Else Kröner-Fresenius Stiftung (2017-A25; 2019_EKMS.23; EKFS Clinician Scientist professorship), and by the DFG–project 404521405, SFB1389—UNITE Glioblastoma, WP B06. M.P. was supported by grants from the DFG–Project ID 394046768 (SFB1366—Vascular Control of Organ Function, WP C01) and the ERC Advanced Grant ‘Characterizing and Harnessing T-cells in the Brain’ (CENTRIC-BRAIN - project 101141901). T.B. was supported by the Medical Faculty Mannheim and the University Hospital Mannheim. L.B was supported by the Dr. Rolf M. Schwiete Foundation, the Swiss Cancer Foundation (Swiss Bridge Award), the Else Kröner Fresenius Foundation (2019_EKMS.49), the University Heidelberg Foundation (Hella Bühler Award), the DFG (German Research Foundation), project 404521405 (SFB1389 UNITE Glioblastoma B03), the Hertie Foundation, and the University of Heidelberg, ExploreTech! Grant.

