## Supplementary data incl methods with method tables and supplementary figures for "Tissue resident CD4+ memory T-cells mark response to immune checkpoint inhibition in high-grade glioma"

### Supplementary Information

This file contains detailed methods, two supplementary method tables, six supplementary figures, and supplementary references to Turco Tan et al., entitled: Tissue resident CD4<sup>+</sup> memory T-cells mark response to immune checkpoint inhibition in high-grade glioma

### Methods

#### *Mice*

Specific and Opportunistic Pathogen free (SOPF) female C57Bl/6J mice were bought from Janvier Laboratories and used at the age of 6-12 weeks. All animal protocols were performed in compliance to the laboratory animal research guidelines and were approved by the governmental authorities (animal protocols: G-27/17, G-130/23, Regional Administrative Authority Karlsruhe, Germany).

#### *Cell lines*

Murine glioblastoma GL261 cells were purchased from the National Cancer Institute Tumor. The murine fibroblast OVA-transfected cell line BOK was kindly provided by Prof. Dr. R. Offringa, German Cancer Research Center (DKFZ), Heidelberg, Germany. All cells were cultured in high glucose Dulbecco's modified Eagle's medium (DMEM, D6429) supplemented with 10 % fetal bovine serum (FBS, F0804) and 100 U/ml penicillin and 100 µg/ml streptomycin (P4333; all Sigma-Aldrich) at 37°C, 5% CO<sub>2</sub>, routinely tested for contamination by multiplex cell contamination test (Multiplexion GmbH), and were not passaged for more than ten times.

#### *Tumor cell inoculation and ICI treatment*

10<sup>5</sup> GL261 murine glioblastoma cells were diluted in 2 µl sterile PBS (Sigma-Aldrich) and stereotactically implanted into the right hemisphere of C57Bl/6J mice (coordinates: 2 mm right lateral of the bregma and 1 mm anterior to the coronal suture with an injection depth of 3mm below the dural surface) using a 10µl Hamilton micro-syringe driven by a fine step stereotactic device (Stoelting). On day 13, allocation to treatment groups was performed based on MRI-based tumor volume (see below). On days 13,

16, and 19, 100 µg per mouse anti-CTLA-4 (9D9, BioXcell) and 250 µg per mouse anti-PD-1 (RMP1-14, BioXcell), or equivalent doses of isotype control antibodies (MCP-11 and 2A3, BioXcell), were administered by intraperitoneal injection.

##### *Peptide vaccination*

C57Bl/6J mice were immunized by subcutaneous injection of 100 µg peptide in Montanide-ISA51 (Seppic) emulsion as described previously<sup>1</sup>. Prior to immunization, peptides were diluted in DMSO and subsequently in PBS for a final DMSO concentration of 10% v/v under sterile conditions. Then, they were emulsified with an equal volume of Montanide-ISA51 adjuvant. Mice were treated with two subcutaneous injections of a peptide-adjuvant emulsion of each 50µl into both lateral pectoral regions. At the first immunization on day 1, mice additionally received 300 ng recombinant murine GM-CSF (PeproTech) by subcutaneous injection, and 5% imiquimod cream (Aldara, Meda Pharma) was applied to the shaved injection sites. Control mice received Montanide emulsion without peptides. Mice were boosted on day 10 with 100 µg peptide in Montanide-ISA51 emulsion and Aldara as before. and splenocytes were isolated on day 21.

##### *MR imaging and tumor response criteria*

Tumor-bearing animals were imaged on day 13, 19 and 26 after intracranial tumor injection using a 9.4 Tesla horizontal bore small animal NMR scanner (BioSpec 94/20 USR, Bruker BioSpin GmbH) with a four-channel phased-array surface receiver coil under isoflurane anesthesia. Tumor volumes were calculated based on T2 weighted images, by manually segmenting the images in the Osirix or ITKsnap imaging software. Classification of response and non-response was based on RANO criteria as previously established (10). All tumor volume and response calculations were done blinded.

##### *Isolation of murine tumor-infiltrating lymphocytes*

For isolation of tumor-infiltrating lymphocytes, mice were cardially perfused after receiving a lethal dose of narcotics. The cerebellum was removed and only the tumor-bearing hemisphere was excised. After enzymatical digestion (HBSS (Sigma Aldrich) with 50 µg/ml Liberase D (Roche)) for 30 min at 37°, the tissue was meshed twice through a 100 µm and 70 µm cell strainer to obtain a single cell suspension. Myelin removal was done with a 30% Percoll gradient (GE Healthcare, Princeton, NJ, USA) according to the manufacturer's instruction.

##### *Patient Treatment, Tumor Digestion and Isolation of Immune Cells*

As described previously<sup>2</sup>, recurrent GBM patients undergoing surgery at the University of California, Los Angeles were treated with standard of care therapies, with some receiving off-label, off-trial neoadjuvant pembrolizumab. All patients provided written informed consent. This study was conducted in accordance with the Declaration of Helsinki, and followed a protocol approved by an institutional review board (UCLA Medical Institutional Review Board 2, IRB#10-000655-AM-00059). CD45+ immune cells from a tumor tissue piece were isolated by magnetic bead positive selection (Miltenyi) according to manufacturer's instructions. CD45+ cell counts and tissue weights were used to calculate per patient immune cell densities. Immune cells were subjected to singel cell RNA Sequencing.

##### *Generation of TCR RNA*

TCR amplification and addition of the T7 promotor was done using primers MmTRAC co mRNA rev and T7 EF1a F (Supplementary Table 1). PCR was set up with PCR Master mix (Takara; Clone Amp HiFi PCR Premix, 639298). Following PCR cycles were used:

|  |  |  |
| --- | --- | --- |
| 98°C | 30 sec | **30 cycles** |
| 98°C | 5 sec |  |
| 55°C | 5 sec |  |
| 72°C | 12 sec |  |
| 72°C | 15 sec |  |

**Supplementary Table 1. Primers**

| Target | Forward 5' – 3' | Reverse 3' – 5' |
| --- | --- | --- |
| MmTRAC_co mRNA_<br>rev | TCAGCTGGACCACAGTCTCAGG | / |
| T7 EF1a F | TAATACGACTCACTATAGGGAC<br>AGAACACAGGCCACCATG | / |

#### *Isolation of human peripheral blood monocyctic cells*

Human peripheral blood monocytic cells (PBMCs) were isolated from research-only buffy-coats from healthy donors provided by Blutspendezentrale IKTZ Heidelberg. Buffy-Coats were anticoagulated using EDTA and stored on ice before processing. Briefly, PBMCs were isolated by density gradient centrifugation using Biocoll Separation Solution (Biochrom). Samples were centrifuged at 800g without brake at room temperature. PBMCs were subsequently used for rapid expansion protocol.

#### *Rapid expansion of PBMCs*

Expansion of PBMC for TCR electroporation was done using irradiated feeder PBMC from three donors. Cells were expanded by stimulation with human anti-CD3 antibody (OKT-3, BioLegend) and human IL-2 (Novartis) in X-Vivo-15 medium (Lonza; BE02-060F) supplemented with 100 U/ml penicillin and 0.1 mg/ml streptomycin (Sigma-Aldrich; P4333), 2.5 µg/mL amphotericin B (ThermoFisher; 15290018), 20 µg/mL Gentamycin (ThermoFisher; 15750060), and 2% human albumin (Sigma-Aldrich, SRP6182) for 14 days, according to the rapid expansion protocol (97) (REP) as described previously<sup>3</sup>. Upon sufficient proliferation, REP-PBMC were split and their media was supplemented with fresh IL-2.

#### *TCR electroporation*

Freshly produced REP-PBMCs were electroporated with TCR-RNA using the program EO-115 for primary cells on the 4D Nucleofector (Lonza) according to the manufacturer's instructions and allowed to rest overnight before being used for TCR testing. REP-PBMCs were then kept at room temperature for 10 minutes to allow cell pores to close. Subsequently, REP-PBMCs were carefully plated in pre-warmed TCR electroporation solution (TexMACS, Miltenyi, 130-097-196; supplemented with 2% human AB serum).

#### *TCR testing*

TCR-transduced expanded T (TCR-T) cells were harvested, resuspended in X-Vivo15 medium (Lonza; BE02-060F) supplemented with 2% human AB serum (Sigma Aldrich) and cell numbers were quantified using Trypan blue. To investigate TCR reactivity against tumor cell lines, GL261, GL261-OVA I, and BOK (overexpressing OVA) cell lines were expanded for 7 to 10 days. Prior assessing TCR reactivity, tumor cells were

pre-treated with 100 ng/mL IFN-g (Peprotech, 315-05) for 24 hours to increase MHC I and II expression. 75,000 tumor cells were plated on top of 150,000 TCR-T-cells for each tested condition. As positive controls, TCR-T-cells were stimulated with 20 ng/ml phorbol myristate acetate (PMA, Sigma-Aldrich) and 1 µg/ml ionomycin (Sigma-Aldrich) (positive control), or CD3/CD28 TransAct Beads (Miltenyi). Unstimulated TCR-T-cells were used as a background control. After setting up the co-culture, 5 µL of anti-human CD107a APC-H7 antibody (BD Biosciences) was supplemented per well. After 1 hour of incubation at 37 °C in 5% CO<sub>2</sub>, cytokines were trapped using GolgiStop, and GolgiPlug (BD Biosciences), following the manufacturer's instructions. Following another 4-hour incubation, cells were washed and stained with flow cytometry antibodies, as described before. Subsequently, cells were analyzed at BD FACS Lyric.

#### *Flow cytometry*

To prevent unspecific antibody binding, murine cells were pre-incubated with anti-mouse CD16/CD32 (eBioscience; clone 93; 14-0161). Human cells were blocked using 10% human AB serum (Sigma-Aldrich; H4522) for 10 min prior to extracellular staining. Surface markers were labeled in FACS buffer (or PBS when fixable viability dyes were used) for 30 min at 4°C. For intracellular cytokine detection, cells were incubated with Brefeldin A (5 µg/ml, Sigma-Aldrich) for 5 hours at 37°C in 5% CO<sub>2</sub> to allow cytokine accumulation. Fixation and permeabilization were performed after surface staining, according to the manufacturer's protocols using the Foxp3/Transcription Factor Staining Buffer Set or the Intracellular Fixation & Permeabilization Buffer Set (both eBioscience), followed by intracellular staining in 1× permeabilization buffer for 45 min at 4°C. Antibody concentrations were optimized by prior titration. Fluorescence minus one (FMO) controls were included in all experiments. A comprehensive list of antibodies used is provided in Supplementary Table 2. Flow cytometry data were acquired using the Lyric Flow Cytometer (BD Biosciences), or Canto II System (BD Biosciences), and analyzed with FlowJo software (versions 9 or 10).

Lymphocytes were subjected to fluorescence-activated cell sorting (FACS) on FACS Aria II (BD Biosciences; Germany) through an 85 mM nozzle and 4-way purity. Cells were sorted into regular 1.5ml tubes (Eppendorf) containing 5µl PBS supplemented with 0,04% bovine serum albumin and kept on ice until processing. Discrimination of dead cells was performed using fixable viability dye eFluor780 (eBioscience). Alive single CD45<sup>high</sup> CD3<sup>+</sup> cells were sorted as T cells from GL261 tumors and used for single cell sequencing.

**Supplementary Table 2. Flow cytometry antibodies and dyes.**

| Target | Label | Clone | Order information |
| --- | --- | --- | --- |
| Fixable viability dye | eFluor780 | / | eBioscience; 65-0865 |
| Fixable viability dye | eFluor506 | / | eBioscience; 65-0866 |
| mCD45 | BV510 | 30-F11 | Biolegend; 103137 |
| mCD11b | APC | M1/70 | Biolegend; 101212 |
| mCD3 | FITC | 17A2 | Biolegend; 100204 |
| mCD4 | Pacific Blue | RM4-5 | Biolegend; 100730 |
| mCD8 | AF700 | 56-6.7 | Biolegend; 100730 |
| H-2Kd | APC | SF1-1.1 | Biolegend; 116619 |
| I-A/I-E | PE-Cy7 | M5/114.15.2 | eBioscience; 25-5321-80 |
| OVA | PE | 25-D1.16 | BioLegend; 141603 |
| hCD3 | BV510 | HIT3A | BD Biosciences; 564713 |
| hCD4 | FITC | SK3 | BD Biosciences; 344604 |
| hCD8 | BV421 | RPA-T8 | BD Biosciences; 562428 |
| mTCRb | PE | H57-597 | BioLegend; 109207 |
| hCD107a | APC-H7 | H4A3 | BD Biosciences; 561343 |
| hTNF-alpha | BV711 | MAb11 | BioLegend; 502939 |

##### *Single Cell RNA sequencing library preparation and sequencing*

*Murine samples.* Single cell RNA sequencing of sorted CD45<sup>+</sup> CD3<sup>+</sup> T-cells from ICB R and NR murine tumors was performed using the 10x genomics V2 3' Kit according to manufacturer protocol. Libraries were sequenced on Illumina HiSeq4000 at a depth of at least 20000 reads per cell as per recommendation.

*Human samples.* Cell preparation, library preparation, and sequencing were carried out according to Chromium product-based manufacturer's protocols (10x Genomics). Sequencing was carried out on a NovaSeq 6000 S2 2 x 50 bp flow cell (Illumina) utilizing either the Chromium single-cell 3' or 5' gene expression library construction per the manufacturer's protocol.

##### *Single Cell RNA Sequencing Analysis*

*Murine data.* Single cell RNA sequencing data was aligned using cellranger (v5.1.2). The resulting count matrices were imported into R and analyzed using Seurat. SoupX was used to remove background noise and miQC was used to remove cells with high mitochondrial expression. Cells with less than 2000 nCount\_RNA and 1000

nFeature\_RNA were discarded. Non-T-cells were excluded from analysis. After quality control, cells were log normalized with a scale factor of 10,000 and integrated using Harmony. 40 harmony components were used for downstream analysis. Cells were first annotated using CD4, CD8, TRDC and TRGC as CD4, CD8 and gamma delta T-cells respectively. Cells were subset into CD4 or CD8 and clustered with graph-based clustering method. Differential gene expression was performed, and top gene expression was used to annotate each cell cluster. Uniform manifold approximation and projection (UMAP) was plotted to visualize cell clusters. Data was imported into Scanpy for umap density and stacked violin plot visualization. EnhancedVolcano was used to visualize differential gene expression as volcano plot.

*Human data.* Raw FASTQ reads were aligned with Cell Ranger version 7.0.1 (10x Genomics) to the Genome Reference Consortium Human Build 38 (GRCh38). Aligned data were analyzed with Seurat, version 4.2.0. Cells with >20% mitochondrial RNA, or fewer than 200 expression genes, and genes expressed in less than 20 cells were discarded. The SCTransform function was used for normalization, scaling, selection of variable features, and regressing out mitochondrial gene expression, ribosomal gene expression, number of transcripts detected, number of unique genes detected, and predicted cell cycle score. All samples were integrated into one object for downstream analysis using a standard Seurat integration pipeline. PCA and UMAP were performed for dimensional reduction of the dataset and for viewing. Unbiased clustering of cells at various resolutions was performed to partition into cell types, with differentially expressed genes being calculated using the FindAllMarkers and FindMarkers functions to confirm cell identity. Identified PTPRC (CD45) expressing immune cells were sub-divided into CD8A or CD8B expressing lymphocytes, CD8A- and CD8B- lymphocytes, and myeloid cells for further analyses. Sub-clustering was performed on these populations by extracting their raw count slots, splitting the object by sample, and re-running the Seurat normalization and integration pipeline. Cell type densities per patient were calculated by multiplying the frequency of each cell type among total immune cells by the overall immune cell density. For the survival analysis, we stratified the cohort of neoadjuvant pembrolizumab-treated GBM patients by their median cell type density value, and survival probabilities were calculated based on the Kaplan-Meier estimator.

#### *Whole exome sequencing*

Whole exome sequencing (WES) of GL261 tumor tissue from ICI responder and non-responder mice was conducted as described previously<sup>4</sup>. Briefly, DNA from GL261 tumor tissue from R and NR mice was extracted using the INVISORB® DNA Tissue Mini Kit (STRATEC Biomedical AG) according to the manufacturer's instruction. Exome sequencing was performed on the Illumina NextSeq500 platform (Illumina Inc, San Diego, Calif.) using High output flow cell (75 nt reads paired end + 8 nt index). SureSelectXT Target Enrichment System (Agilent Technologies) was used for library generation according to the manufacturer's instructions.

#### *Neoepitope MHC-binding prediction*

MHC binding prediction of potential GL261-associated neo-epitopes identified by WES was performed using NetMHC 4.0<sup>5</sup> (MHC class I, H2-D<sup>b</sup> and H2-K<sup>b</sup>), and NetMHCII 2.3<sup>6</sup> (MHC class II, H2-IA<sup>b</sup>). 27-mer peptides containing the central amino acid substitution were used as input. Only predicted epitopes containing the amino acid substitution were selected for further analyses. Binder strength thresholds followed the NetMHC server guidelines.

#### *ELISpot (Enzyme-linked-immuno-Spot)*

5 x 10<sup>5</sup> splenocytes from vaccinated mice were seeded onto 96-well ELISpot plates (MabTech, MAIPSWU10) that had been coated with IFN-gamma-capture antibody according to manufacturer's instructions (MabTech, 3321-2A). Splenocytes were stimulated for 36h with the respective peptide of interest, MOG (35-55) (control peptide), 0.1% DMSO (vehicle control), and 20 ng ml<sup>-1</sup> phorbol myristate acetate (PMA, Sigma-Aldrich) with 1 µg ml<sup>-1</sup> ionomycin (Sigma-Aldrich) (positive control) at 37 °C and 5% CO<sub>2</sub>. To determine MHC restriction, peptide-pulsed splenocytes were incubated with purified blocking antibodies against murine MHC class I (28-8-6, BioLegend) or MHC class II (M5/114.15.2, BioLegend) during in vitro stimulation prior to IFN-γ ELISpot analysis. IFN-gamma producing T-cell populations were analyzed using mouse IFN-gamma ELISPOT assay (MabTech, 3321-2A) according to the manufacturer's instructions and quantified by an Immunospot Analyzer (Cellular Technology Ltd). T-cell IFN-gamma responses were defined by the mean spot numbers per 5x10<sup>5</sup> effector cells. Three technical and at least three biological replicates were used for ELISpot experiments.

### Supplementary Figures

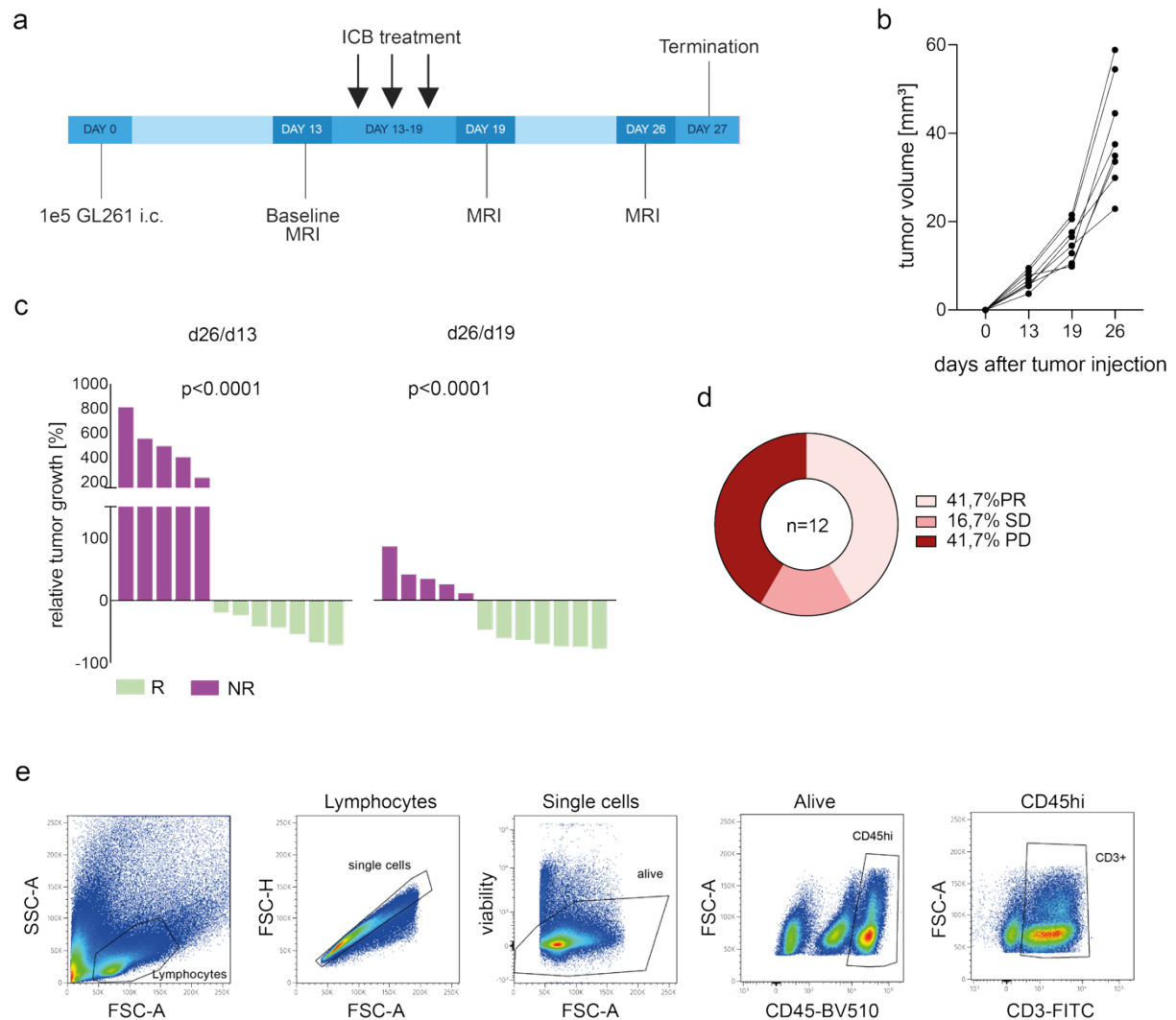

Figure S1

**Figure S1: Tumor growth dynamics define ICI treatment response in GL261-bearing mice.** **a**, Experimental workflow. C57Bl/6J mice were treated repeatedly with 250 µg anti-PD-1 and 100 µg anti-CTLA-4 (ICI) on d13, d16 and d19 after intracranial GL261 tumor inoculation. Tumors were excised on d27 for analysis of tumor-infiltrating T-cells. **b**, Quantified tumor growth in control treated mice (n=8). **c**, Waterfall plots depicting relative tumor growth throughout the experiment between baseline MRI (d13) and final MRI (d26, left) and during the late phase between post-treatment MRI (d19) and final MRI (d26, right). N(R, light green) = 7, n(NR, purple) = 5. T-test, two-tailed. **d**, Donut plot of PR, SD, and PD frequencies of ICI treated mice by iRANO criteria. N(all ICI treated) = 12, n(PR) = 5, n(SD) = 2, n(PD) = 5. R is defined as PR and SD;

NR is defined as PD. PR, partial response; SD, stable disease; PD, progressive disease. **e**, Gating strategy for FACS of CD3<sup>+</sup> T-cells from ICI-treated GL261 tumors as in (a). CD45<sup>hi</sup>, highly CD45 positive cells. Related to Fig. 1.

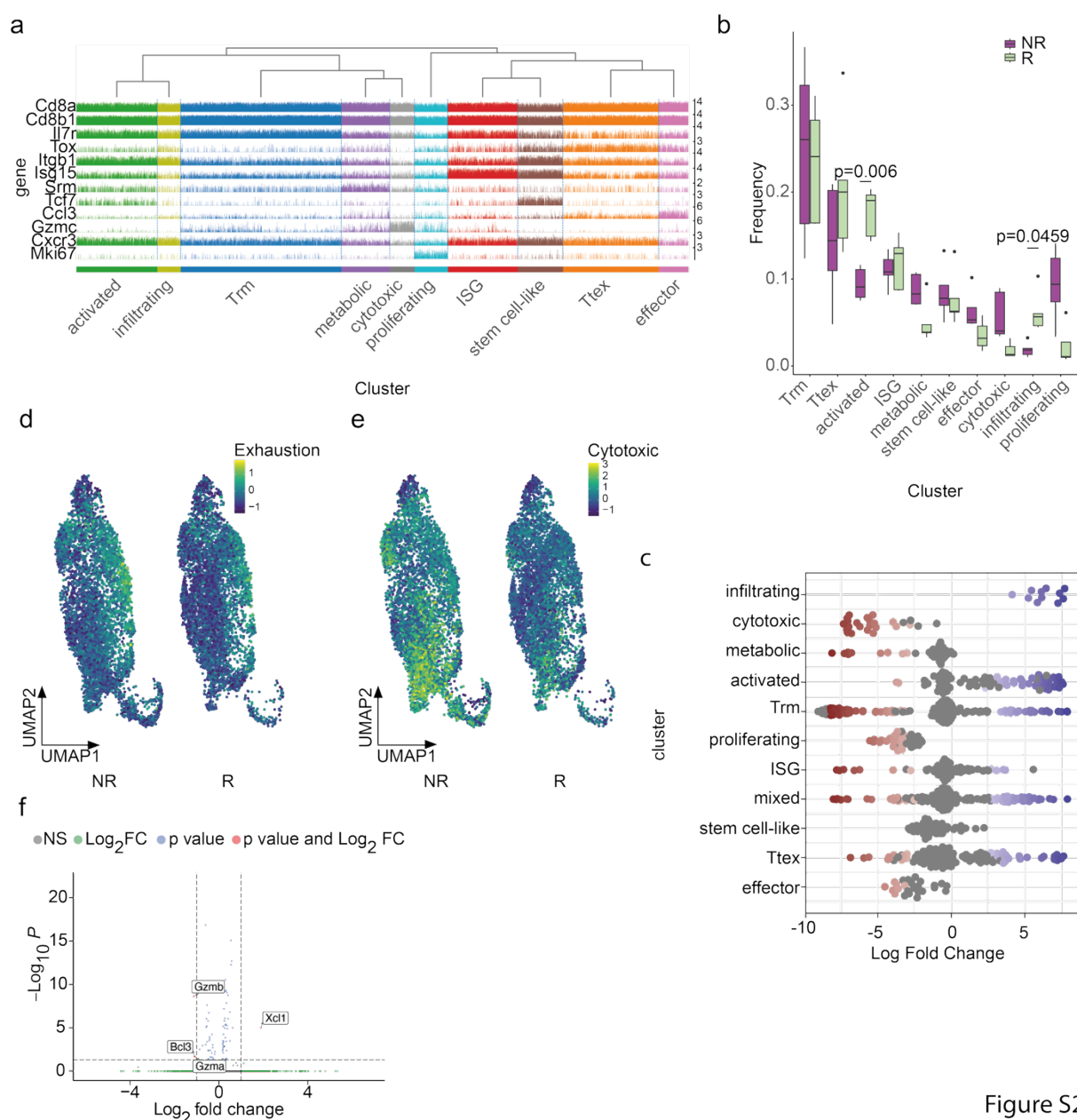

Figure S2

**Figure S2: CD8<sup>+</sup> T-cell transcriptomic clusters differ in ICI R and NR GL261 tumors.** **a**, Trackplot of cluster-defining genes in CD8<sup>+</sup> T-cells from ICI R and NR. **b**, Frequencies of CD8<sup>+</sup> T-cell clusters from ICI R (light green, n=5) and NR (purple, n=5) mice. **c**, Relative CD8<sup>+</sup> T-cell cluster abundance in ICI R compared to NR mice as in (b). Only significant p values are depicted. Statistical significance was determined by t-test with Holm correction. **d,e**, Exhaustion (d) and cytotoxicity (e) scores based on the expression of *Pdcd1*, *Ctla4*, *Havcr2*, *Lag3*, *Tigit*, *Tox* (d), and *Prf1*, *Gzma*, *Gzmb*, *Gzmk*, *Ngk7* (e), respectively, projected onto CD8<sup>+</sup> T-cell UMAP from ICI NR (left) and R (right). **f**, Volcano plot showing differentially expressed genes (DEG) in CD8<sup>+</sup> *Tcf7*<sup>+</sup> stem cell-like T-cell cluster in ICI R vs. NR mice. Only significant differences are labelled. Statistical significance was determined by Wilcoxon rank sum test with Bonferroni correction. Related to Fig. 2.

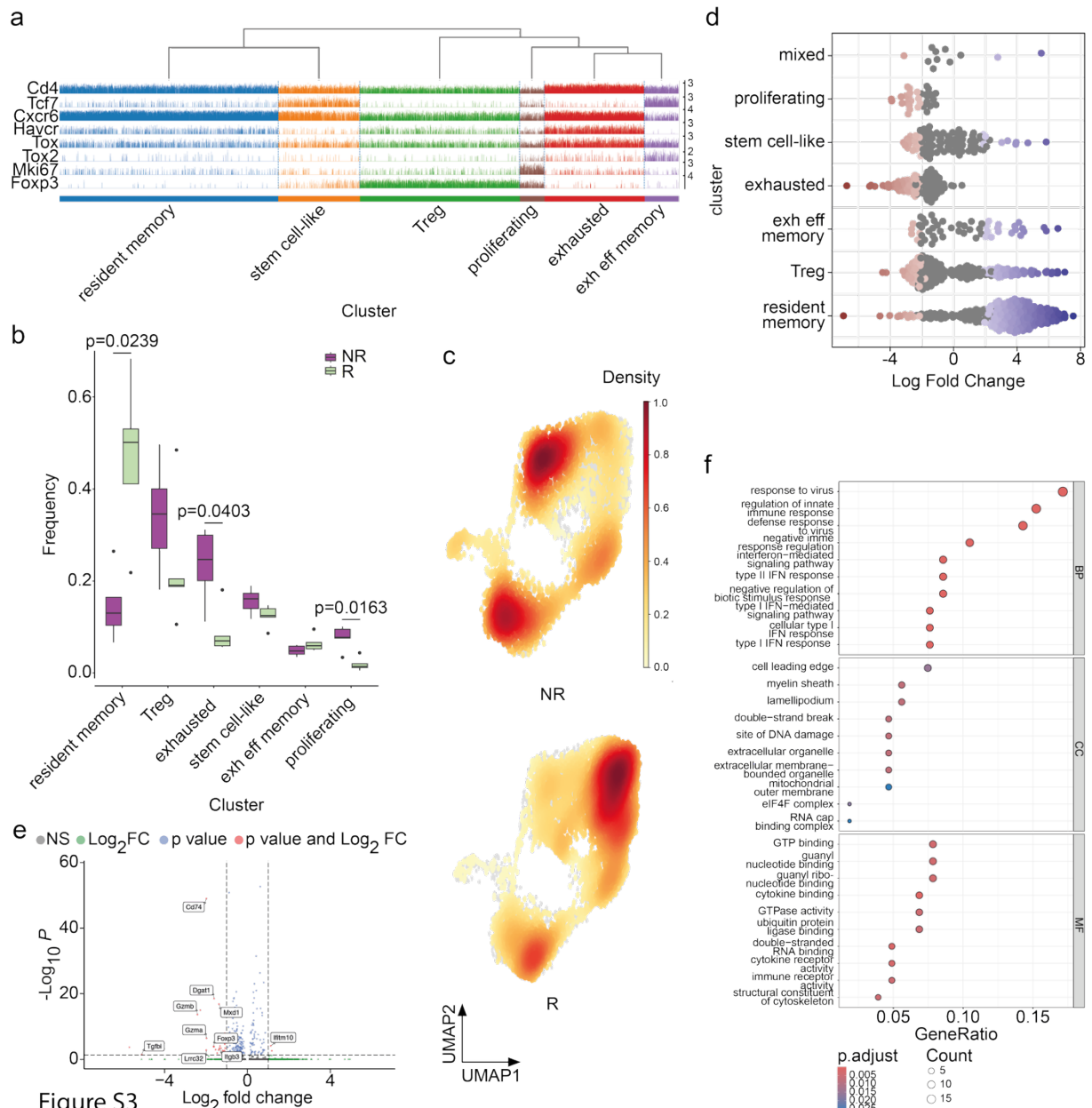

**Figure S3: CD4<sup>+</sup> T-cell transcriptomic clusters differ in ICI R and NR GL261 tumors.** **a**, Trackplot of cluster-defining genes in CD4<sup>+</sup> T-cells from ICI R and NR. **b**, Frequencies of CD4<sup>+</sup> T-cell clusters from ICI R (light green, n = 5) and NR (purple, n = 5) mice. **c**, Density plot showing distribution of CD4<sup>+</sup> T-cell clusters in ICI NR and R mice projected onto CD4<sup>+</sup> T-cell UMAP from ICI NR (top) and R (bottom). **d**, Relative CD4<sup>+</sup> T-cell cluster abundance in ICI R compared to NR mice as in (b). **e**, Volcano plot showing differentially expressed genes (DEG) in CD4<sup>+</sup> *Tcf7*<sup>+</sup> stem cell-like T-cell cluster in ICI R vs. NR mice. Only significant differences are labelled. Wilcoxon rank sum test with Bonferroni correction. **f**, Gene ontology analysis of DGE in CD4<sup>+</sup> T-cells from R and NR. Statistical significance was determined by one-sided Fisher's Exact Test with Benjamini-Hochberg correction. Related to Fig. 2.

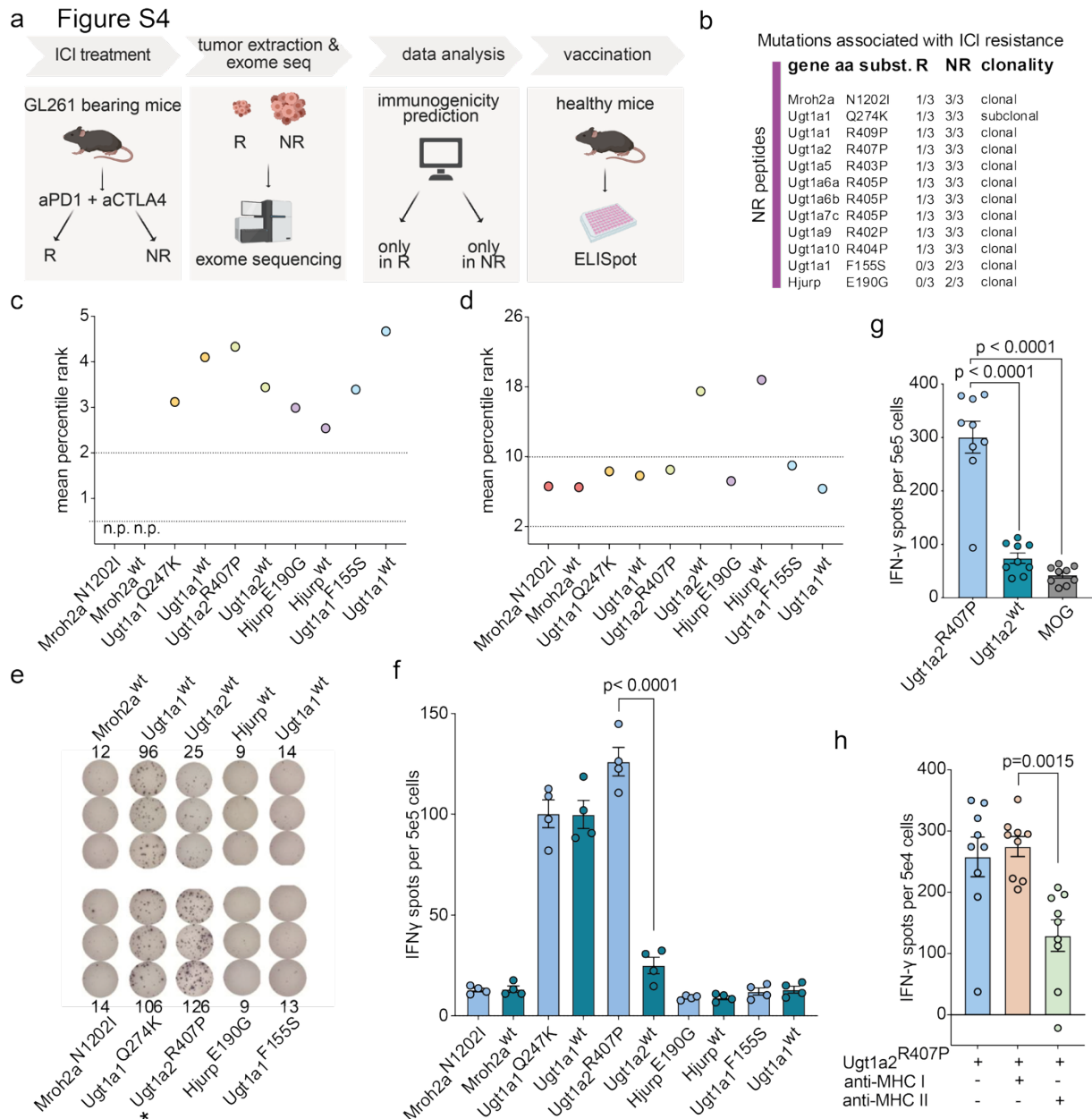

**Figure S4: Discovery of GL261 neoepitopes using ICI R and NR mice.** **a**, Workflow for sample processing, exome-sequencing, analysis, and neoepitope immunogenicity testing. Mice were treated and tumors excised as described in Fig S1. Tumor samples were subjected to exome sequencing to derive mutated gene expression of tumor cells. Immunogenicity of mutations was in silico predicted and tested via vaccination of mice and IFN-γ ELISpot of splenocytes. **b**, Overview of mutations found in ICI NR tumors. N(R) = 3, n(NR) = 3. aa subst., amino acid substitution. **c,d**, Mean percentile immunogenicity ranks of neoepitopes from (b) and their respective wildtype counterparts by in silico immunogenicity prediction for C57BL/6 alleles H2-D<sup>b</sup> and H2-k<sup>b</sup> (c), and H2-IA<sup>b</sup> (d) using NetMHC. Mean percentile rank <0.5, strong binder; mean percentile rank >0.5 and <2.0, weak binder; mean percentile rank >2.0, no binder, for H2-D<sup>b</sup> and H2-k<sup>b</sup> in (c). Mean percentile rank <2, strong binder; mean percentile rank

>2 and <10, weak binder; mean percentile rank >10, no binder, for H2-IA<sup>b</sup> in (d). wt, wildtype; n.p., no prediction. **e**, IFN-γ release of splenocytes measured by ELISpot after vaccination of mice with the respective mutated peptide and *ex vivo* recall with indicated mutated and corresponding wt peptides. Representative images with mean spot numbers for each peptide. N = 4 mice per peptide. Asterisk marks mutation-specific immunogenic peptide. **f**, Quantification of (e). Mean MOG spot number subtracted. **g,h**, IFN-γ release of splenocytes measured by ELISpot after vaccination of mice with neoepitope Ugt1a2(R407P) (n = 9 mice) and *ex vivo* recall with indicated peptides (f), or with Ugt1a2(R407P) and MHC class I or class II blocking antibodies (g). Mean MOG spot number subtracted in (h). MOG, myelin oligodendrocyte protein epitope, negative control. Mean ± s.e.m. (e,g,h). One-way ANOVA (e,g,h) with Tukey's test for multiple comparison for (g). Related to Fig. 3.

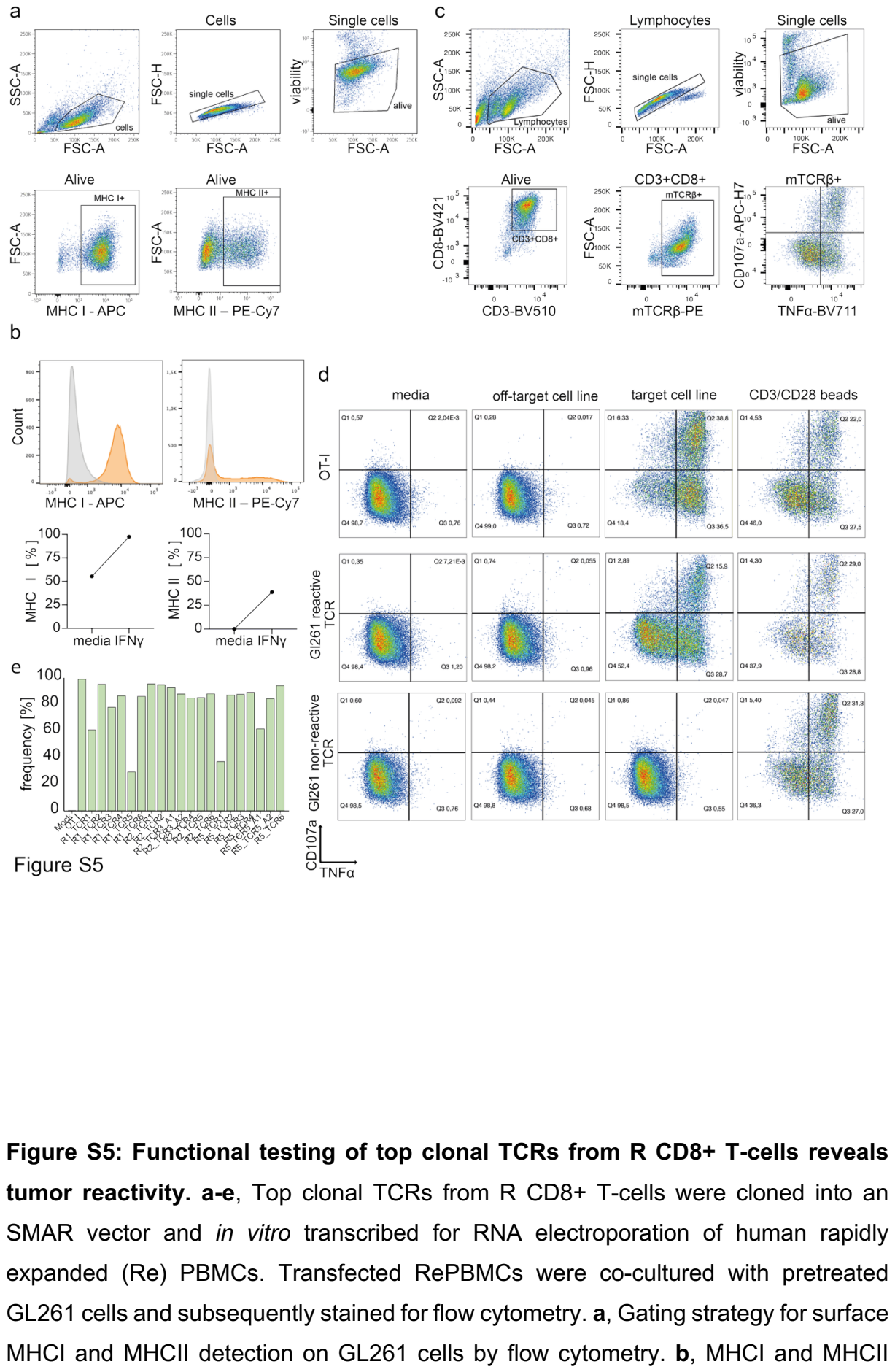

Figure S5: Functional testing of top clonal TCRs from R CD8+ T-cells reveals tumor reactivity. a-e, Top clonal TCRs from R CD8+ T-cells were cloned into an SMAR vector and in vitro transcribed for RNA electroporation of human rapidly expanded (Re) PBMCs. Transfected RePBMCs were co-cultured with pretreated GL261 cells and subsequently stained for flow cytometry. a, Gating strategy for surface MHCI and MHCII detection on GL261 cells by flow cytometry. b, MHCI and MHCII

expression of GL261 cells pretreated with IFN-γ *in vitro*, measured by flow cytometry and analyzed as shown in (a). Top, histograms. Orange, treated; grey, Isotype control. Bottom, quantified frequency of MHC positive cells. Media, untreated. Representative of at least 2 independent experiments. **c**, Gating strategy for surface transgenic TCRβ and CD107a and intracellular TNFα detection on TCR-transfected RePBMCs by flow cytometry. **d**, CD107a surface and intracellular TNFα levels of TCR-transfected TCRβ<sup>+</sup> T-cells co-cultured with GL261 cells (targetT-cell line) or OT-1-expressing BOK cells (off-targetT-cell line), measured by flow cytometry and analyzed as shown in (c). Representative reactive TCRs are shown. OT-I TCR was used as control; here, GL261 represented off-targetT-cell line and BOK represented targetT-cell line. **e**, Surface transgenic TCRβ expression of TCR-transfected RePBMC T-cells, measured by flow cytometry and analyzed as shown in (d). Frequency of all alive cells is shown. Related to Fig. 3.

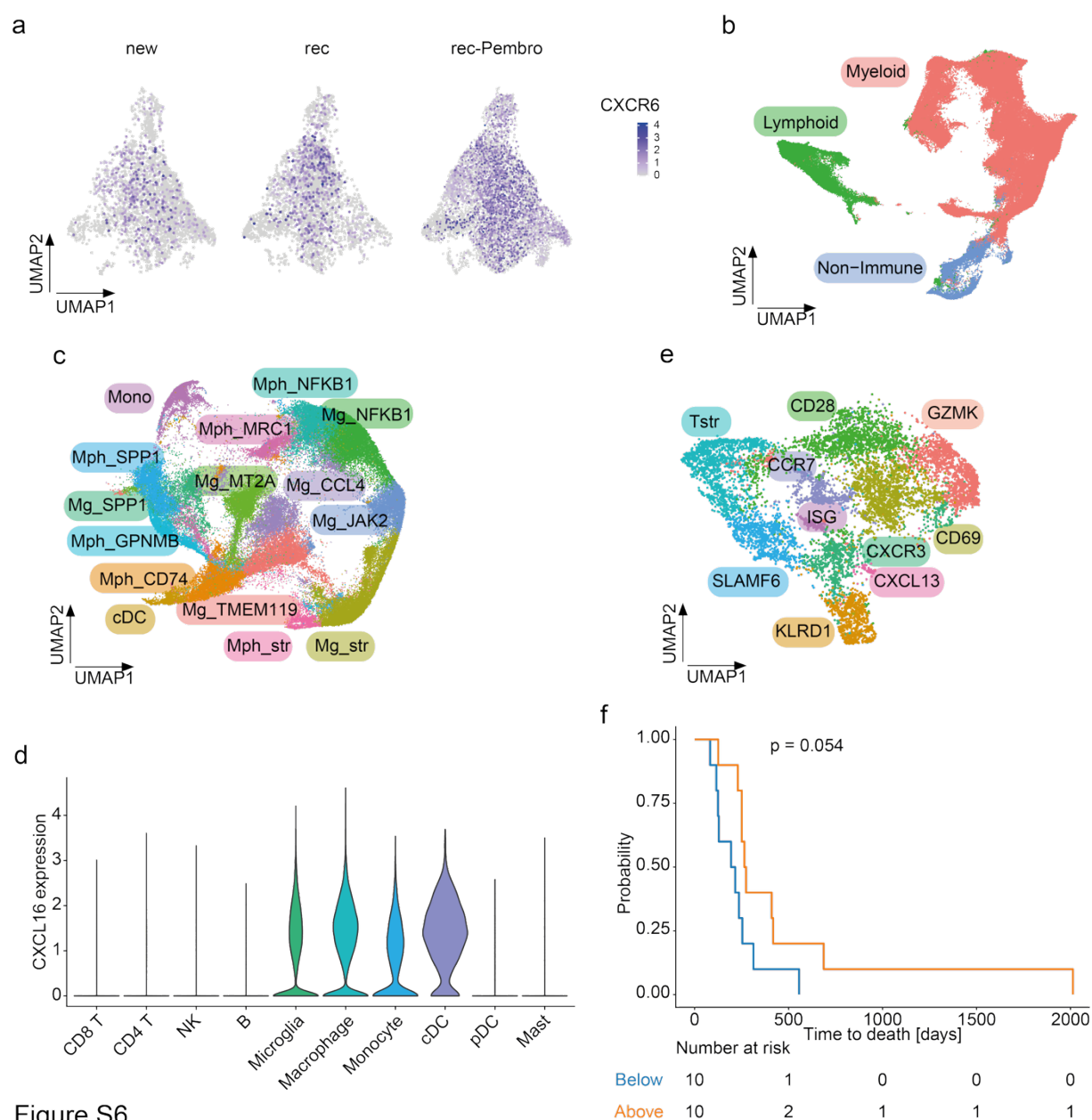

Figure S6

**Figure S6: CXCR6 expression in intratumoral T-cells from recurrent glioblastoma patient samples treated with neoadjuvant pembrolizumab.** **a**, CXCR6 expression levels in T-cells from newly diagnosed (new), recurrent (rec), and pembrolizumab-treated recurrent (rec-Pembro) glioblastoma samples<sup>27</sup> projected on the respective UMAP. n(new) = 61,269 cells, n(rec) = 38,683 cells, n(rec-Pembro) = 56,814 cells. **b-g**, Patients presenting with recurrent glioblastoma were treated with neoadjuvant pembrolizumab (n=20) or left untreated (n=15) in addition to standard of care temozolomide before surgical resection. A tissue fragment was weighed and used for magnetic bead enrichment of CD45+ immune cells, which were subjected to single cell RNA sequencing. **b**, UMAP of all sorted CD45+ cells. Myeloid cluster annotation based on *PTPRC* (encoding CD45) and *CD14* expression. Lymphoid cluster annotation based on *PTPRC* expression and *CD14*-. Non-immune, *PTPRC*- *CD14*-

cells. Phenotypic clusters are represented in distinct colors. N = 136,407 cells. **c**, UMAP of myeloid cells from (b). Transcriptomic clusters are represented in distinct colors. N = 83,790 cells. **d**, *CXCL16* expression levels by CD45<sup>+</sup> cell clusters from (b) in all tumor samples (n=35 tumors and 97,232 cells). **e**, UMAP of *CD8A*<sup>+</sup> or *CD8B*<sup>+</sup> lymphoid cells from (b), indicating CD8<sup>+</sup> T-cell clusters. Transcriptomic clusters are represented in distinct colors. N = 5,933 cells. **f**, Kaplan-Meier plot of overall survival probability of glioblastoma patients treated with neoadjuvant pembrolizumab according to *CXCR3*<sup>+</sup> T-cell density among CD8<sup>+</sup> T-cells. Orange, above median *CXCR3*<sup>+</sup> T-cell density; blue, below median *CXCR3*<sup>+</sup> T-cell density. N(above) = 10, n(below) = 10. Mantel-Cox log-rank test. Related to Fig. 4.
